# Kinesin-binding protein remodels the kinesin motor to prevent microtubule-binding

**DOI:** 10.1101/2021.06.02.446814

**Authors:** April L. Solon, Zhenyu Tan, Katherine L. Schutt, Lauren Jepsen, Sarah E. Haynes, Alexey I. Nesvizhskii, David Sept, Jason Stumpff, Ryoma Ohi, Michael A. Cianfrocco

**Affiliations:** Department of Cell and Developmental Biology, University of Michigan; Department of Biophysics, University of Michigan; Life Sciences Institute, University of Michigan; Department of Molecular Physiology and Biophysics, University of Vermont; Department of Biomedical Engineering, University of Michigan; Department of Pathology, University of Michigan; Department of Computational Medicine and Bioinformatics, University of Michigan; Department of Biological Chemistry, University of Michigan

**Author notes:** These authors contributed equally to this work (April L. Solon, Zhenyu Tan). For correspondence &.

## Abstract

Kinesins are tightly regulated in space and time to control their activation in the absence of cargo-binding. Kinesin-binding protein (KIFBP) was recently discovered to bind the catalytic motor heads of 8 of the 45 known kinesin superfamily members and inhibit binding to microtubules. In humans, mutation of KIFBP gives rise to Goldberg-Shprintzen syndrome (GOSHS), but the kinesin(s) that is misregulated to produce clinical features of the disease is not known. Understanding the structural mechanism by which KIFBP selects its kinesin binding partners will be key to unlocking this knowledge. Using a combination of cryo-electron microscopy and crosslinking mass spectrometry, we determined structures of KIFBP alone and in complex with two mitotic kinesins, revealing regions of KIFBP that participate in complex formation. KIFBP adopts an alpha-helical solenoid structure composed of TPR repeats. We find that KIFBP uses a 2-pronged mechanism to remodel kinesin motors and block microtubule-binding. First, KIFBP engages the microtubule-binding interface and sterically blocks interaction with microtubules. Second, KIFBP induces allosteric conformational changes to the kinesin motor head that displace a key structural element in the kinesin motor head (α-helix 4) required for microtubule binding. We identified two regions of KIFBP necessary for *in vitro* kinesin-binding as well as cellular regulation during mitosis. Taken together, this work establishes the mechanism of kinesin inhibition by KIFBP and provides the first example of motor domain remodeling as a means to abrogate kinesin activity.

## INTRODUCTION

Kinesins comprise a superfamily of microtubule-based motor proteins that play essential roles in virtually every aspect of cell physiology, including mitotic spindle assembly, regulation of microtubule dynamics, ciliogenesis, and transportation of cargoes throughout the cell^1–4^. A signature protein fold shared among all members of the kinesin superfamily is a catalytic “motor” domain. The kinesin motor domain contains binding sites for both microtubules and ATP, enabling these proteins to convert energy from ATP hydrolysis into mechanical force^5^. In most kinesin motors, this catalytic cycle powers motility of the proteins along microtubule tracks. While the motor domain exhibits structural and high sequence conservation among the superfamily, sequence differences imbue each kinesin with unique characteristics and are responsible for diversifying motor functions within the cell. In addition, the non-motor regions of different kinesin family members have diverged to confer specificity for cargo binding and regulation.

Kinesins are regulated at many levels to ensure that they become activated at the right time and place. Auto-inhibition, wherein kinesins adopt a conformation that prevents microtubule-binding^6–8^; sequestration within the nucleus^9–14^; and cell cycle-dependent protein expression^15^ are common strategies to prevent untimely motor-track interactions. Kinesins are also regulated by post-translational modifications, *e*.*g*., phosphorylation, which can serve to activate microtubule-binding^16–18^. Lastly, kinesin-interacting proteins such as adaptors and light chains, and their phosphorylation, can regulate the ability of transport kinesins to engage cargo^1,19–24^, or target them to specific locations within the cell^25,26^. KIFBP, a new class of kinesin-binding protein, has emerged as an important negative regulator of a subset of kinesin motors^27,28^.

KIFBP was discovered as a disease-causing gene associated with the neurological disorder Goldberg-Shprintzen syndrome (GOSHS^29–31^), an autosomal disease characterized by facial dysmorphism, mental retardation, and congenital heart disease (OMIM #609460). In mice and zebrafish, loss of KIFBP function leads to neuronal migration and maturation defects in the developing brain^32,33^. Emerging data demonstrate a compelling role for KIFBP in regulating motor-microtubule interactions for 8 of the 45 kinesin motors encoded by the human genome. KIFBP interacts directly with the motor head of Kinesin-2 (KIF3A), Kinesin-3 (KIF1A, KIF1B, KIF1C, KIF13B, and KIF14), Kinesin-8 (KIF18A), and Kinesin-12 (KIF15) family members, resulting in inhibition of motor-microtubule binding both *in vitro* and in cells^27,28^. How the regulation of kinesin motors by KIFBP is linked to specific biological processes is largely unexplored, although neuronal microtubule dynamics appear to be controlled through KIFBP-dependent regulation of KIF18A^27^. Moreover, recent work has shown that KIFBP is critical for ensuring proper mitotic spindle assembly by regulating the mitotic kinesins KIF15 and KIF18A^28^.

To understand the mechanism of KIFBP:motor head interaction, we used cryo-electron microscopy (cryo-EM) coupled with crosslinking mass spectrometry (XL-MS) to determine the structure of KIFBP alone and in complex with KIF15 and KIF18A. We show that KIFBP is a tandem repeat protein constructed of 9 helix pairs that assemble into a solenoid-like structure. When complexed with KIF15 and KIF18A, three helices in KIFBP (HP4a/b-HP5) associate closely with the kinesin α4 helix, an interaction that requires a 15 Å displacement of α4 from its resting position. Using molecular dynamics simulations, we find that kinesin α4 is immobile when a motor head is not bound to KIFBP, suggesting that allosteric changes drive the repositioning of α4 required for binding HP4a/b-HP5. Using information obtained from cryo-EM and XL-MS, we identified two regions in KIFBP that are responsible for KIFBP:kinesin interactions *in vitro*, and show that mutations in these regions disable the ability of KIFBP to regulate KIF15 and KIF18A during mitosis. Collectively, our work demonstrates that KIFBP inhibits KIF15 and KIF18A by binding to and stabilizing a conformation of the kinesin motor head that is incompatible with microtubule binding.

## RESULTS

### KIFBP adopts a solenoid structure composed of TPR motifs

Previous work suggested that KIFBP possessed multiple TPR domains based on primary sequence analysis^27^. Despite this information, the overall structure of KIFBP remained unknown, given that there are no structures of KIFBP fragments or close structural homologs. To understand the overall architecture of KIFBP, we utilized cryo-EM to determine the structure of KIFBP (**Figure 1A, Figure 1 - Supplements 1 & 2, Table 1**). Reconstructions of the full KIFBP molecule at 4.6Å showed that KIFBP is almost entirely alpha-helical, possessing nine helical pairs along with one long helix throughout the 621 amino acid sequence. The alpha helices are arranged into a right-handed superhelical twist, giving KIFBP an appearance analogous to other TPR proteins^34^.

**Figure 1.**
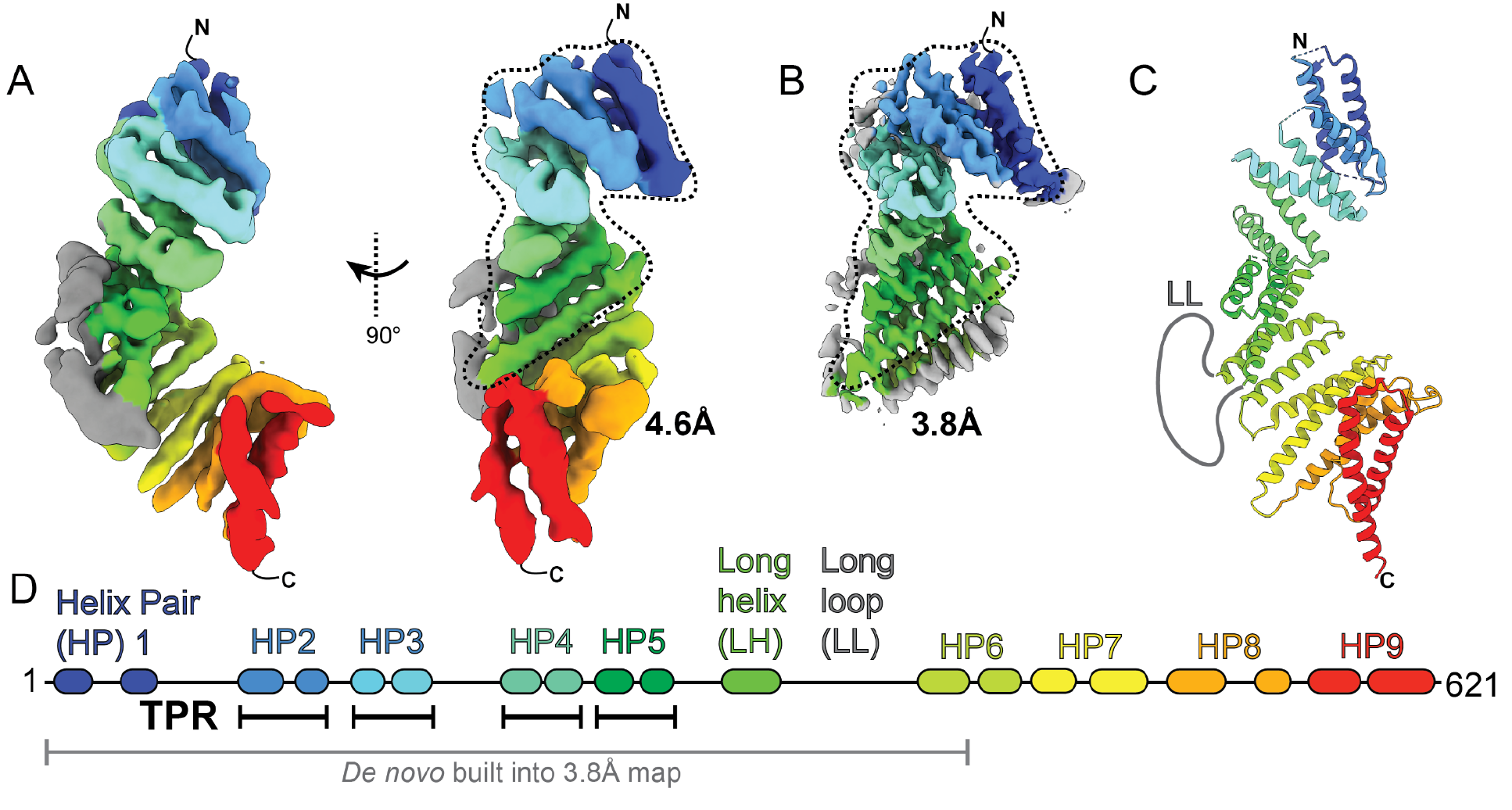
KIFBP adopts a solenoid structure composed of TPR motifs. (A) Overview of KIFBP structure at 4.6Å. Dotted lines indicate the masked region used for the higher-resolution KIFBP core. (B) Structure of KIFBP core at 3.8Å. (C) Combined atomic model of KIFBP. (D) Structural features and nomenclature for KIFBP.

To improve the resolution of KIFBP and enable *de novo* atomic model building, we performed masking and 3D classification on the N-terminal two-thirds of KIFBP to obtain a higher-resolution structure at 3.8Å (**Figure 1B**). We could unambiguously identify amino acid side chains at this resolution, allowing us to construct an atomic model for amino acids 5 to 403 (**Figure 1C, Figure 1 - Supplements 1, 3, 4, and 5, Table 2**). Our atomic model provides a high-confidence positioning of KIFBP residues, allowing us to map the structure onto the KIFBP sequence (**Figure 1D**), confirming alpha-helical positions and showing the locations of loops connecting helical pairs throughout the structure.

### KIFBP inhibits KIF15 microtubule-binding through rearrangement of KIF15’s motor domain

After determining the structure of KIFBP alone, we used cryo-EM to determine the structure of KIFBP bound to KIF15. To prepare cryo-EM samples, we incubated the purified KIF15 motor domain (amino acids 1-375) with KIFBP and subjected the sample to size exclusion chromatography in the presence of ATP (**Figure 2 - Supplement 1**). SDS-PAGE analysis determined that KIF15 co-migrated with KIFBP in a 1:1 complex, and fractions containing the complex were utilized for cryo-EM sample preparation.

We used cryo-EM to determine a ∼4.8Å resolution structure of KIFBP bound to KIF15 (**Figure 2A, Figure 2 - Supplements 1 & 2, Table 3**). At this resolution, we could unambiguously identify the regions of density corresponding to KIFBP in addition to the motor domain of KIF15 (**Figure 2A**). Unexpectedly, when we docked the structure of KIF15 into the reconstruction, we noticed that the α4 helix of KIF15 was missing. Instead, we noticed the presence of an additional alpha-helical density within a cleft of KIFBP, suggesting the displacement of KIF15-α4 into this cleft during complex formation (**Figure 2B**). In the structure, we see that KIFBP occupies the microtubule-binding surface of KIF15, sterically blocking access to the microtubules by KIF15.

**Figure 2.**
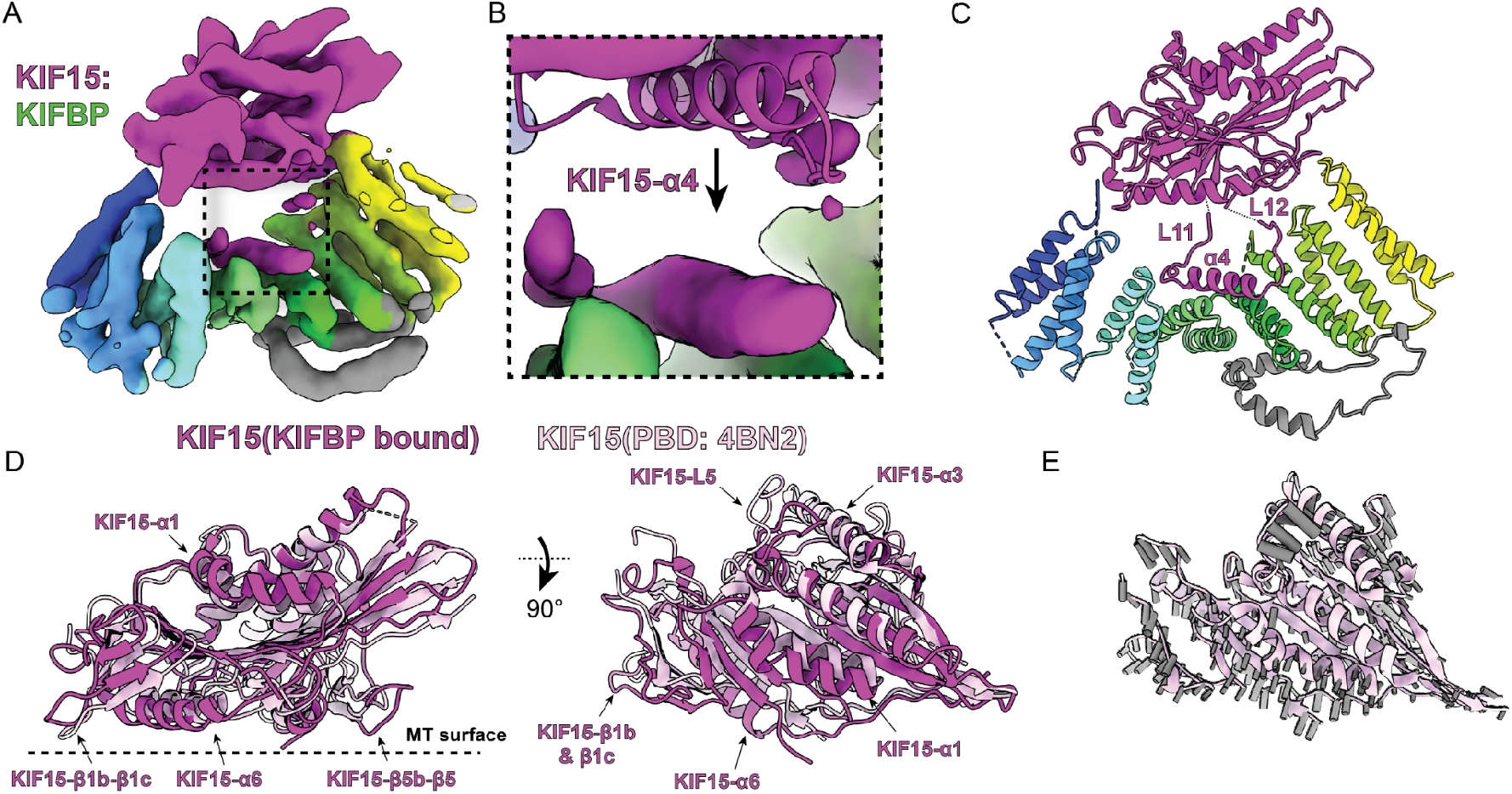
KIFBP stabilizes KIF15 in a conformation that blocks microtubule-binding. (A) KIF15:KIFBP reconstruction. (B) Zoom-in on additional density present in KIF15:KIFBP reconstruction (purple) alongside docked crystal structure (PDB: 4BN2)^36^. (C) Atomic model of KIF15:KIFBP. (D) Superposition of KIF15 bound to KIFBP (purple) with apo-KIF15 (PDB: 4BN2)^36^ relative to KIF15-a2. Structural elements that differ are indicated by arrows. (E) Vectors (gray) calculated from Ca differences between KIF15(KIFBP-bound) vs. KIF15 superimposed on the apo KIF15 crystal structure (PDB: 4BN2)^36^.

To understand how KIFBP affected the overall architecture of KIF15, we utilized both manual building and Rosetta comparative modeling^35^ to develop a model for the KIFBP-engaged KIF15 motor (**Figure 2C**). Our analysis of KIFBP:KIF15 revealed structural rearrangements of the KIF15 motor by KIFBP to disrupt KIF15’s microtubule-binding interface. The most significant structural change involved the repositioning of KIF15-α4 away from the kinesin motor domain, placing KIF15-α4 15Å away from the location found in the crystal structure of KIF15^36^. Notably, kinesin motors require the α4 helix for engaging microtubules during the kinesin mechanochemical cycle^37^. The adjoining loops on each side of KIF15-α4, Loop-11 (“KIF15-L11”) and Loop-12 (“KIF15-L12”), accommodated the repositioning of KIF15-α4 by facilitating the extension of KIF15-α4 away from the body of the motor domain (**Figure 2C**). Whereas KIF15-L12 remains extended in solution, the KIF15-L11 is positioned away from the motor domain and binds along KIFBP-HP4a.

In addition to seeing changes in KIF15-L11, KIF15-α4, and KIF15-L12, we saw that the overall structure of KIF15 adopted a more open conformation (**Figure 2D & E**). The structure showed the shift of alpha-helices KIF15-α1, -α3, and -α6 away from the core of the motor. We observed large movements for beta-strand pairs KIF15-β1b-β1c and KIF15-β5b-β5 in addition to loop KIF15-L5. These changes indicate that KIFBP stabilizes several structural changes in KIF15 to block microtubule-binding. Thus, KIFBP blocks microtubule binding by sterically preventing microtubule interaction in addition to allosterically altering the KIF15 motor.

### KIFBP utilizes a hydrophobic cleft to bind KIF15-α4

Given that KIFBP binds along the microtubule-binding interface of KIF15, we sought to compare the interaction interface between KIF15:KIFBP and KIF15:αβ-tubulin^38^. First, we noticed that the length of the α4 helix is shorter for KIF15-KIFBP compared to the microtubule-engaged α4 helix (**Figure 3A-F**). The length of α4 in KIFBP:KIF15 is similar to the crystal structure of KIF15 when not bound to microtubules^36^. Second, we see that KIF15-L11 is bent relative to α4 at an angle of ∼120° (**Figure 3B**), whereas KIF15-L11 on the microtubule adopts a helical structure to extend the length of α4 (**Figure 3E**)^38^. These two observations indicate that KIFBP holds KIF15-α4 in a conformation that is incompatible with microtubule-binding.

**Figure 3.**
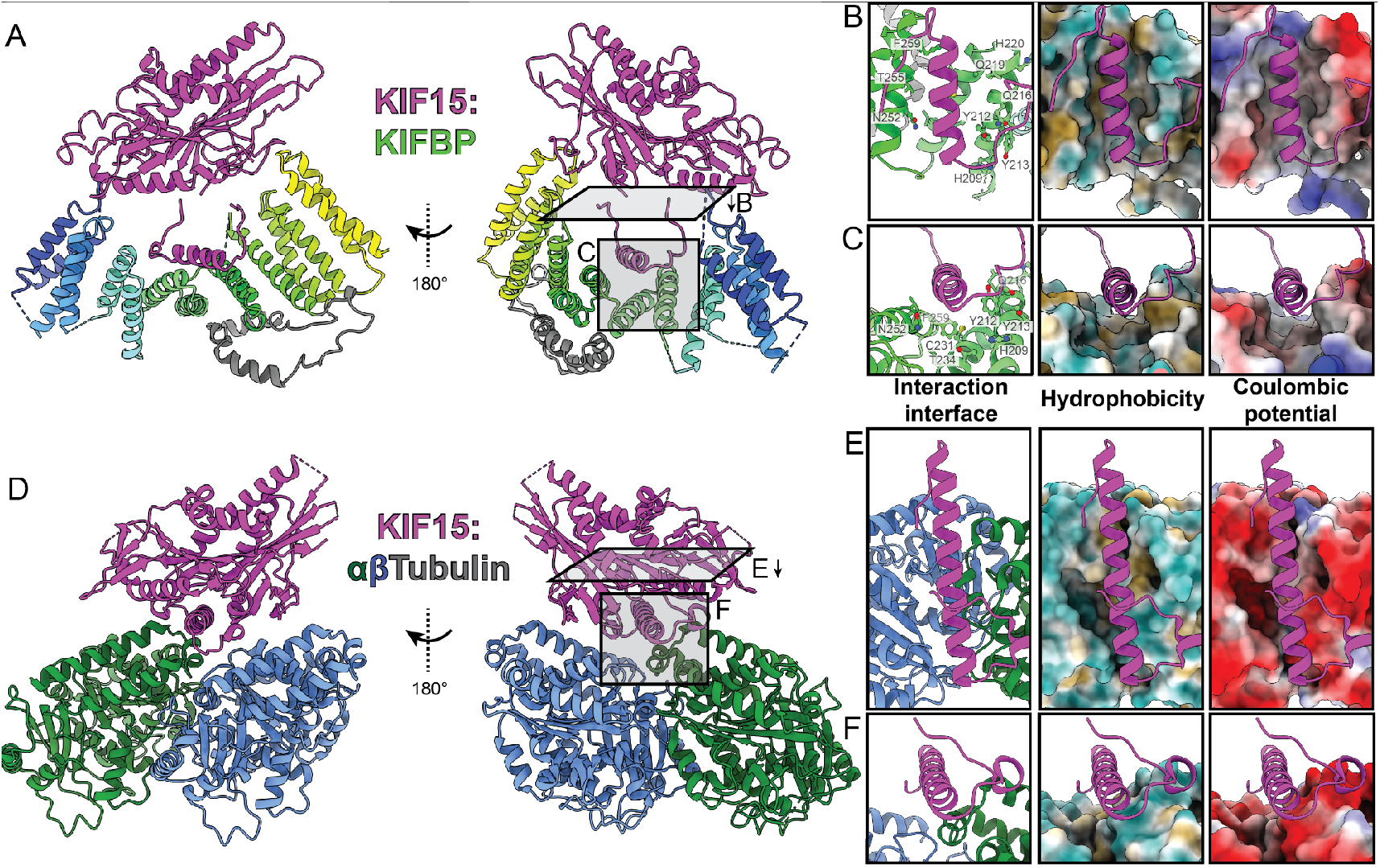
KIFBP utilizes a hydrophobic cleft to bind KIF15-α4. Comparison of KIFBP- and α β-tubulin-bound KIF15. (A) KIF15:KIFBP atomic model. (Right) Gray rectangles indicate viewing directions for panels (B) & (C). (B) & (C) Top and side views of KIF15-α4 interface (left), hydrophobicity (center), and Coulombic potential (right). (D) KIF15: α β-tubulin structure (PDB: 6ZPI)^38^ shown relative to KIF15:KIFBP (A). (Right) Gray rectangles indicate viewing directions for (E) & (F). (E) & (F) Top and side views of KIF15-α4 interface (left), hydrophobicity (center), and Coulombic potential (right). Hydrophobicity lipophilic potential scale: -20 (green, hydrophilic) to 20 (brown, hydrophobic). Coulombic electrostatic potential scale: -10 (red) to 10 (blue) kcal/(mol e).

Comparing the hydrophobicity and electrostatic charge surfaces on KIFBP vs. αβ-tubulin shows that KIFBP binds KIF15-α4 via hydrophobic groove (**Figure 3B, C, E, & F**). The strong electrostatic nature of αβ-tubulin results in minimal hydrophobic residues contributing to KIF15 binding. Unlike αβ-tubulin, KIFBP utilizes a composite binding site stretching across three helices to bind both hydrophobic and polar residues to interact with KIF15-α4, such as KIFBP-H209, -Y212, -Y213, and -T255. Comparing the overall hydrophobicity and charge distribution indicates that KIFBP binds KIF15-α4 in a manner distinct from αβ-tubulin.

### KIFBP engages the microtubule-binding interface of KIF15 using multiple contact points

To obtain further insight into the regions of KIFBP and KIF15 that interact with each other, we performed crosslinking mass spectrometry (XL-MS). Recombinant KIFBP and KIF15 (1-375) were incubated with the 11 Å lysine-targeting crosslinker BS3, digested with trypsin, and analyzed using tandem mass spectrometry. We identified crosslinked peptides using pLink software (see Materials and Methods). We present all high-confidence crosslinks between KIFBP and KIF15 peptides (e-value >0.05) in **Table 5**, and have displayed them on the primary and secondary structures of KIFBP:KIF15 as well (**Figure 4A & 4B**).

**Figure 4.**
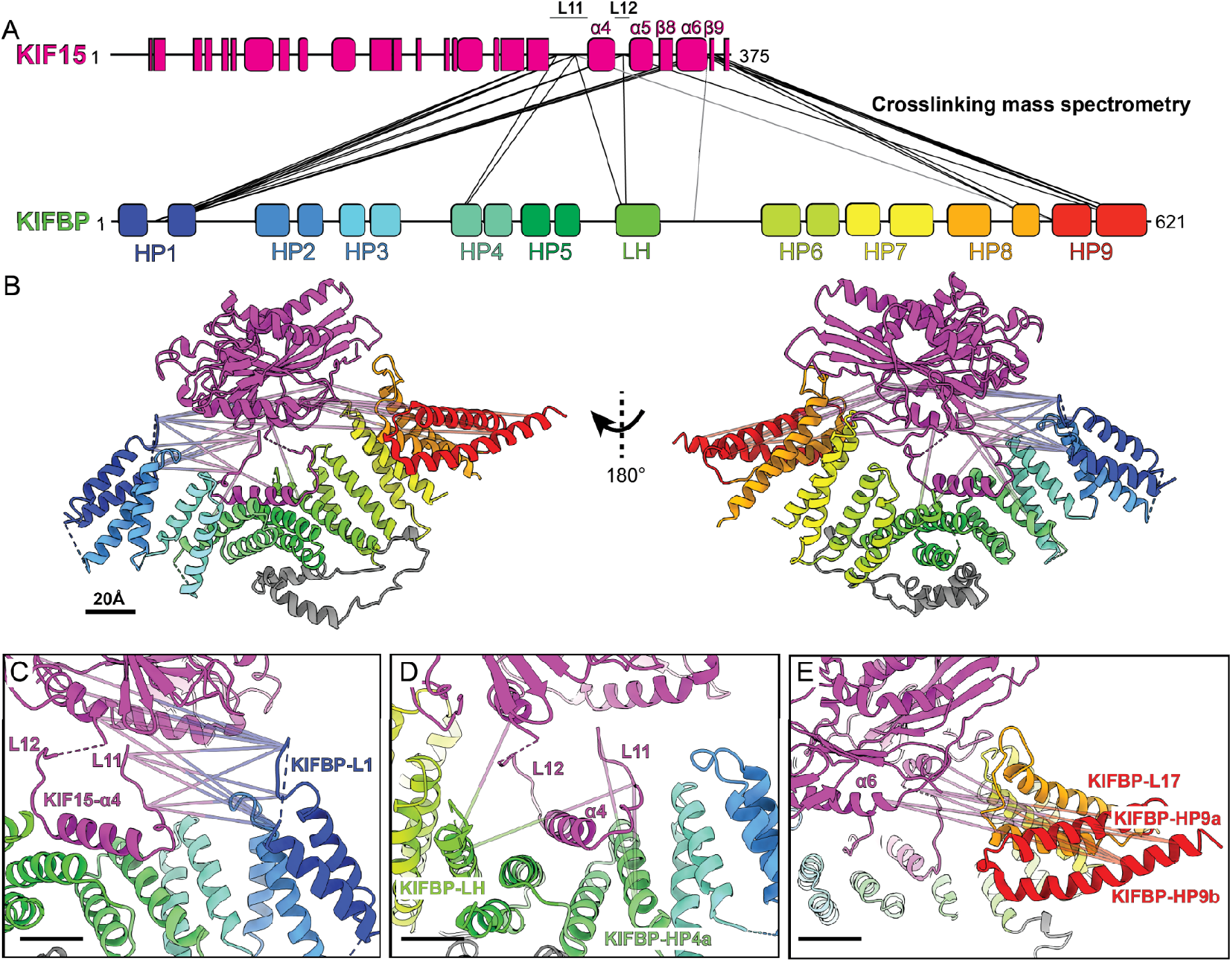
KIFBP physically contacts multiple sites along the microtubule-binding interface of KIF15. (A) Schematic representing the location of identified crosslinks between the KIF15 motor domain (top) and KIFBP (bottom). Secondary structure elements of the two proteins are represented by rectangles (β-sheets), rounded rectangles (α-helices), and lines (unstructured regions). (B) Crosslinks shown in panel A have been superimposed on the cryo-EM structure of KIFBP:KIF15. (C) A zoomed-in view of the crosslinks between KIFBP-L1 and KIF15. (D) A zoomed-in view of the crosslinks between KIFBP-HP4a and -LH and KIF15. (E) Azoomed-in view of the crosslinks between KIFBP-L17, -HP9a, and -HP9b and KIF15. Scale bars are 20Å.

We observed the highest density of crosslinks between three residues of KIFBP-L1 (K26, K30, and K36) and the microtubule-binding interface of KIF15 (K273, K283, K319, and K361) (**Figure 4A & 4C**). Interestingly, these same KIF15 residues also crosslinked to regions in the middle of KIFBP (HP4a, LH, and LL) (**Figure 4D**) and towards the C-terminus (L17, HP9a, and HP9b) (**Figure 4E**). Two KIF15 residues that crosslinked multiple KIFBP sites (K273 and K283) are located in KIF15-L11, adjacent to KIF15-α4. Additionally, residues KIF15-K273 and KIF15-K319 form the kinesin motor microtubule-binding interface^38^. The high density of crosslinks involving these KIF15 residues supports a mechanism of inhibition where KIFBP directly binds the microtubule-binding domain of kinesins, occluding interactions with the microtubule lattice.

KIFBP:KIF15 crosslinks span nearly the entire length of KIFBP to bind the KIF15 microtubule-binding surface. When superimposed onto the KIFBP:KIF15 structure (**Figure 4B-E**), these crosslinks bridge distances greater than the 11 Å BS3 can reach. This suggests that the crosslinked regions of KIFBP may associate transiently with the microtubule-binding interface of KIF15 at different time points during complex formation (see Discussion).

### KIFBP inhibits KIF18A *via* a similar mechanism as KIF15

After characterizing how KIFBP inhibits KIF15 (kinesin-12 family), we next aimed to establish whether KIFBP utilizes the same mode of inhibition for a kinesin motor from a different kinesin family, KIF18A (kinesin-8 family). To determine how KIFBP inhibits KIF18A, we first purified recombinant KIF18A (1-363) motor domain, incubated KIF18A with KIFBP, and performed size exclusion chromatography to confirm the formation of a 1:1 complex (**Figure 5 - Supplement 1**). After preparing cryo-EM grids with the complex, we obtained 2D class averages that appeared similar in shape and features as seen previously for KIF15 (**Figure 5 - Supplement 2**), further confirming the formation of a KIFBP:KIF18A complex.

After performing further single particle analysis, the cryo-EM structure of KIFBP:KIF18A revealed that KIFBP binds KIF18A like KIF15 (**Figure 5A & 5B, Table 4**). In the structure, we see that KIFBP N- and C-terminal domains engage both sides of the motor domain while KIF18A-α4 is displaced away from the motor into the central cavity of KIFBP. To highlight the similarities between KIFBP engagement of KIF15 and KIF18A, we segmented the motor density from either KIFBP:KIF18A (**Figure 5C & 5D**) or KIFBP:KIF15 (**Figure 5E & 5F**). This comparison shows for both motors that 1) α4 is held within the central cleft of KIFBP, 2) Loops-11 & -12 are extended away from the motor, and 3) KIFBP-L11 adopts a curved shape as it makes a ∼120° turn to follow helix KIFBP-HP4a within the KIFBP cleft. Thus, KIFBP stabilization of kinesin α4 helix away from the motor is a shared mode of kinesin inhibition by KIFBP for KIF18A and KIF15.

**Figure 5.**
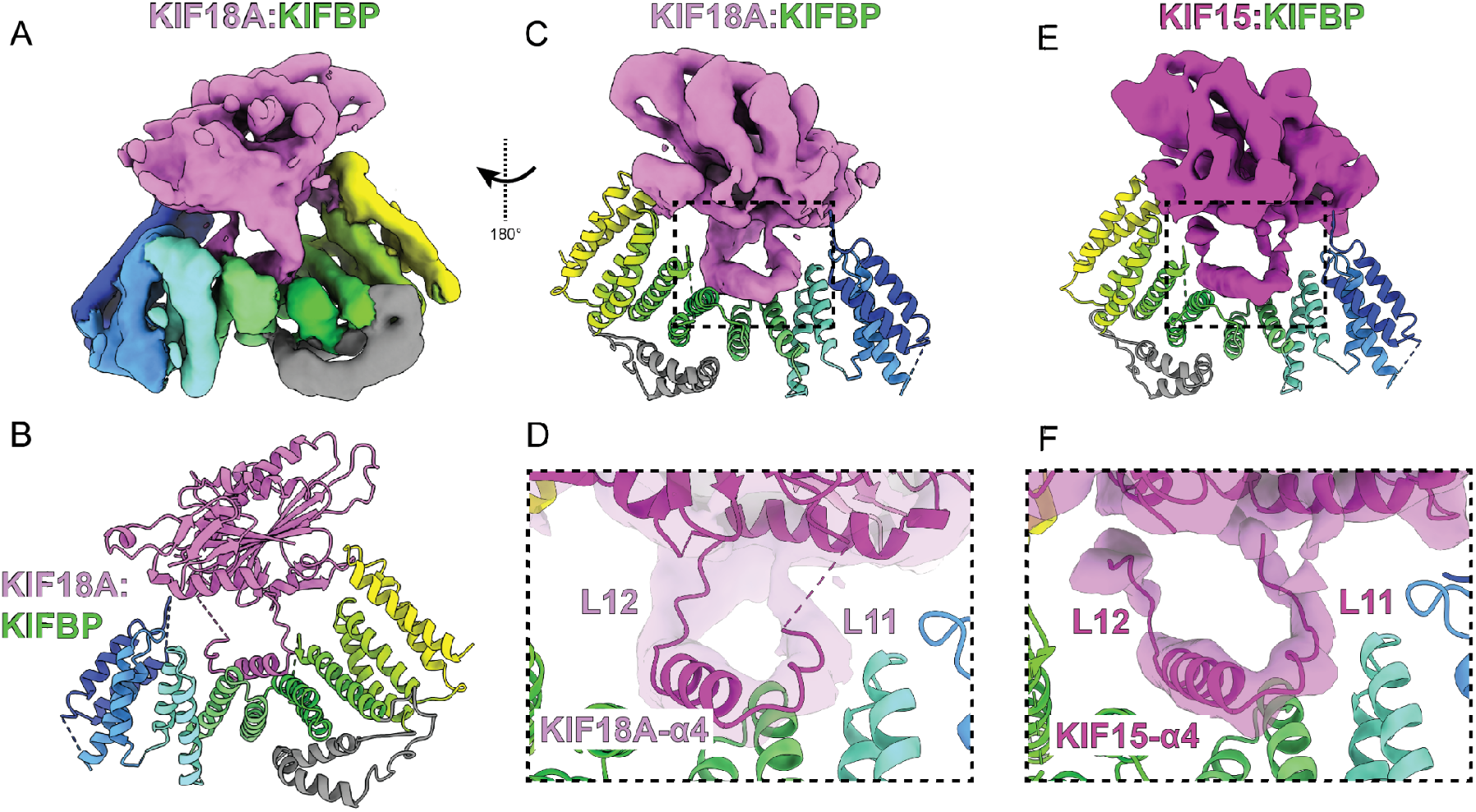
KIFBP inhibits KIF18A *via* a similar mechanism as KIF15. (A) Cryo-EM reconstruction of KIF18A:KIFBP. (B) Atomic model of KIF18A:KIFBP. (C) Segmented KIF18A density shown alongside KIFBP atomic model, rotated 180° from (A). (D) Zoomed-in view of KIF18A density and model on KIFBP interface. (E) Segmented KIF15 density shown in a similar orientation as (C). (F) Zoomed-in view of KIF15:KIFBP interface.

### KIFBP utilizes Loop-1 and Loop-14 to bind kinesin *in vitro*

Our structural data and XL-MS results identified multiple KIFBP:motor interactions that may be important for robust binding and inhibition of motor activity. In particular, our XL-MS results revealed that the most extensively crosslinked residues in KIFBP occurred within Loop-1 and the C-terminal helix pairs of KIFBP (**Figure 4A, Table 5**). Our cryo-EM structure of KIFBP complexed with the motor domains of both KIF15 and KIF18A support a prominent role for Loop-1 in motor engagement. We therefore targeted both KIFBP-L1 and KIFBP-HP9b for mutagenesis, selecting positively and negatively charged residues in those regions and substituting them with alanine or glycine residues (**Figure 6B**). We also constructed a third KIFBP mutant by mutating residues 460-465 in KIFBP-L14 to alanine (**Figure 6B**) because of the proximity of KIFBP-L14 to KIF15 and previous work indicating that it is important for kinesin motor binding^38^.

**Figure 6.**
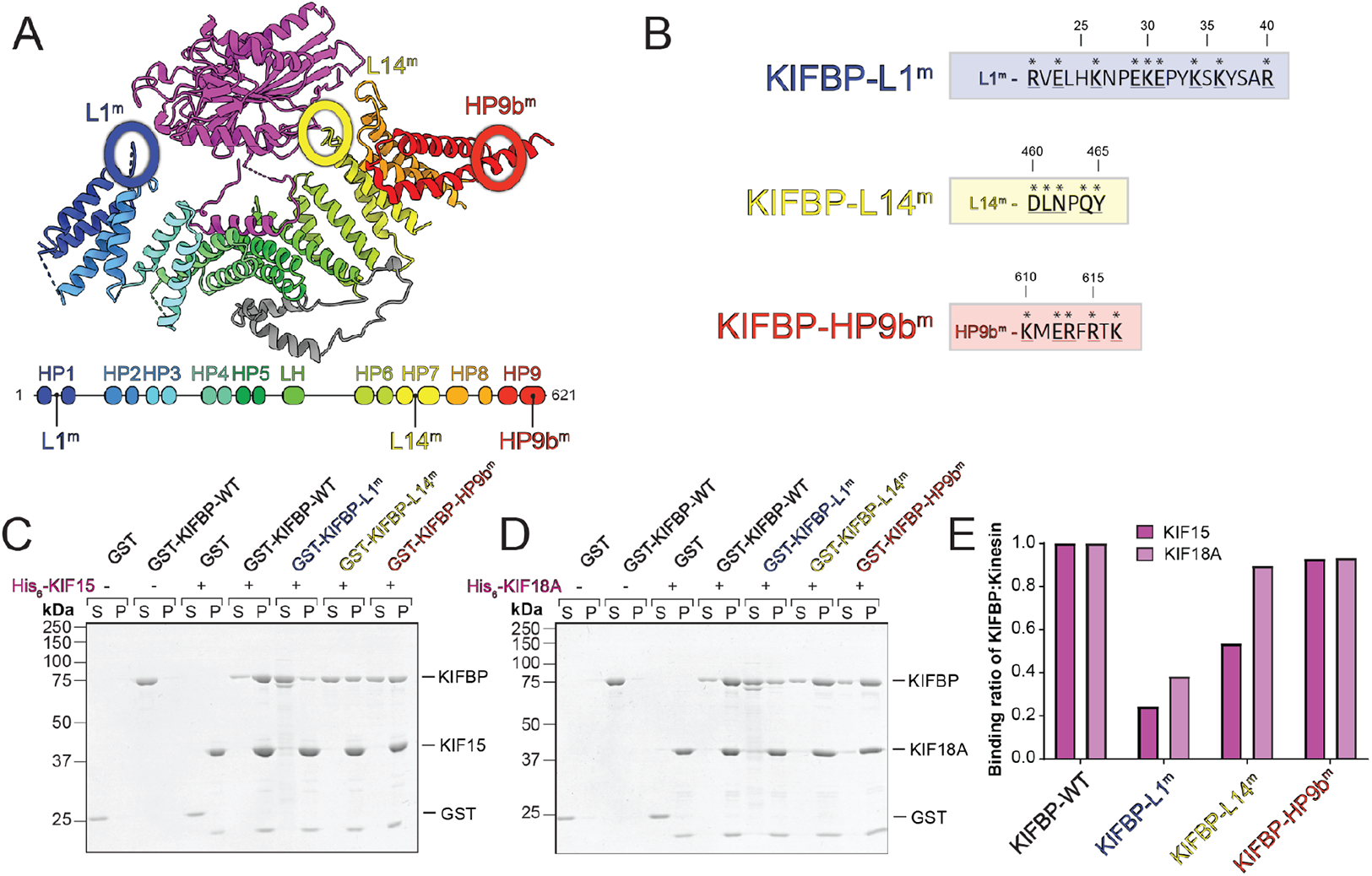
KIFBP utilizes Loop-1 and Loop-14 to bind kinesins *in vitro*. (A) Schematic showing the locations of the three mutations on the cryo-EM structure of KIFBP:KIF15 (top) and on the secondary structure of KIFBP (bottom). Mutations were made to Loop-1, Loop-14, and HP9b of KIFBP. (B) Mutated residues in each mutant are indicated with asterisks; residue position is indicated above each sequence. (C and D) Representative Coomassie gels showing results of in vitro pull-down assays where KIFBP proteins were pulled down by either KIF15 (C) or KIF18A(D). Supernatant and pellet samples, as well as the presence or absence of KIF15 or KIF18A, are indicated at the top of each gel. (E) Quantifications of pull-down assays shown in panels C and D are graphed as the ratio of each KIFBP protein to KIF15or KIF18Ainthe pellet.

First, we tested the ability of these three KIFBP mutants to bind the motor domains of KIF15 and KIF18A *in vitro*. We performed *in vitro* pull-down assays using hexahistidine-tagged motor domains of either KIF15 (1-375) or KIF18A (1-363) immobilized on nickel resin, and analyzed the ratio of recombinant wildtype or mutant KIFBP to kinesin motor domain in the pellet. We used GST as a negative control, which showed little to no nonspecific interaction with nickel resin or immobilized kinesin. When incubated with KIF15, we observed a four-fold reduction in binding of the KIFBP-L1^m^ mutant and a two-fold reduction in binding of the KIFBP-L14^m^ mutant compared to KIFBP-WT, indicating that mutations in these regions abrogate the ability of KIFBP to interact with the motor robustly. Surprisingly, the KIFBP-HP9b^m^ showed no difference in binding compared to KIFBP-WT, suggesting that the charged residues mutated are not essential for motor binding. Thus, although both Loop-1 and the C-terminus of KIFBP were implicated as potentially important for binding in our XL-MS experiment (**Figure 4C & 4E**), mutations to only one of these regions, Loop-1, affected binding biochemically.

Next, we repeated pull-down assays with KIF18A immobilized on the resin and analyzed binding of the same panel of mutants. Similar to KIF15, the KIFBP-L1^m^ mutant showed a three-fold reduction in binding compared to KIFBP-WT. Intriguingly, the KIFBP-L14^m^ mutant showed only a slight 10% reduction in binding to KIF18A compared to KIFBP-WT, in contrast to the 50% decrease in binding to KIF15. Based on these results, Loop-1 may represent a common binding site among all KIFBP-binding kinesins, whereas Loop-14 may be important only for some kinesins including KIF15 but not KIF18A.

### Mutations in KIFBP-L1 and KIFBP-L14 disrupt the regulation of mitotic kinesins

Overexpression of KIFBP in HeLa cells leads to defects in chromosome alignment and an increase in spindle length^28^. To determine whether mutations that block KIFBP interaction *in vitro* also reduce KIFBP effects during mitosis, we transfected N-terminally mCherry-tagged KIFBP constructs into HeLa Kyoto cells and measured chromosome alignment and spindle length in metaphase arrested cells. Consistent with previous results, overexpression of mCherry-KIFBP-WT decreased chromosome alignment, quantified by an increase in full-width at half maximum (FWHM) of centromere fluorescence distribution along the length of the spindle (**Figure 7A & 7B**)^28,39^. Overexpression of mCherry-KIFBP-WT also increased spindle lengths, as shown previously^28^ (**Figure 7A & 7B**). These mitotic effects scaled with the mCherry-KIFBP-WT expression level, where cells expressing higher levels of mCherry-KIFBP-WT had longer spindles and more severe chromosome alignment defects (**Figure 7 - Supplement 1**).

**Figure 7.**
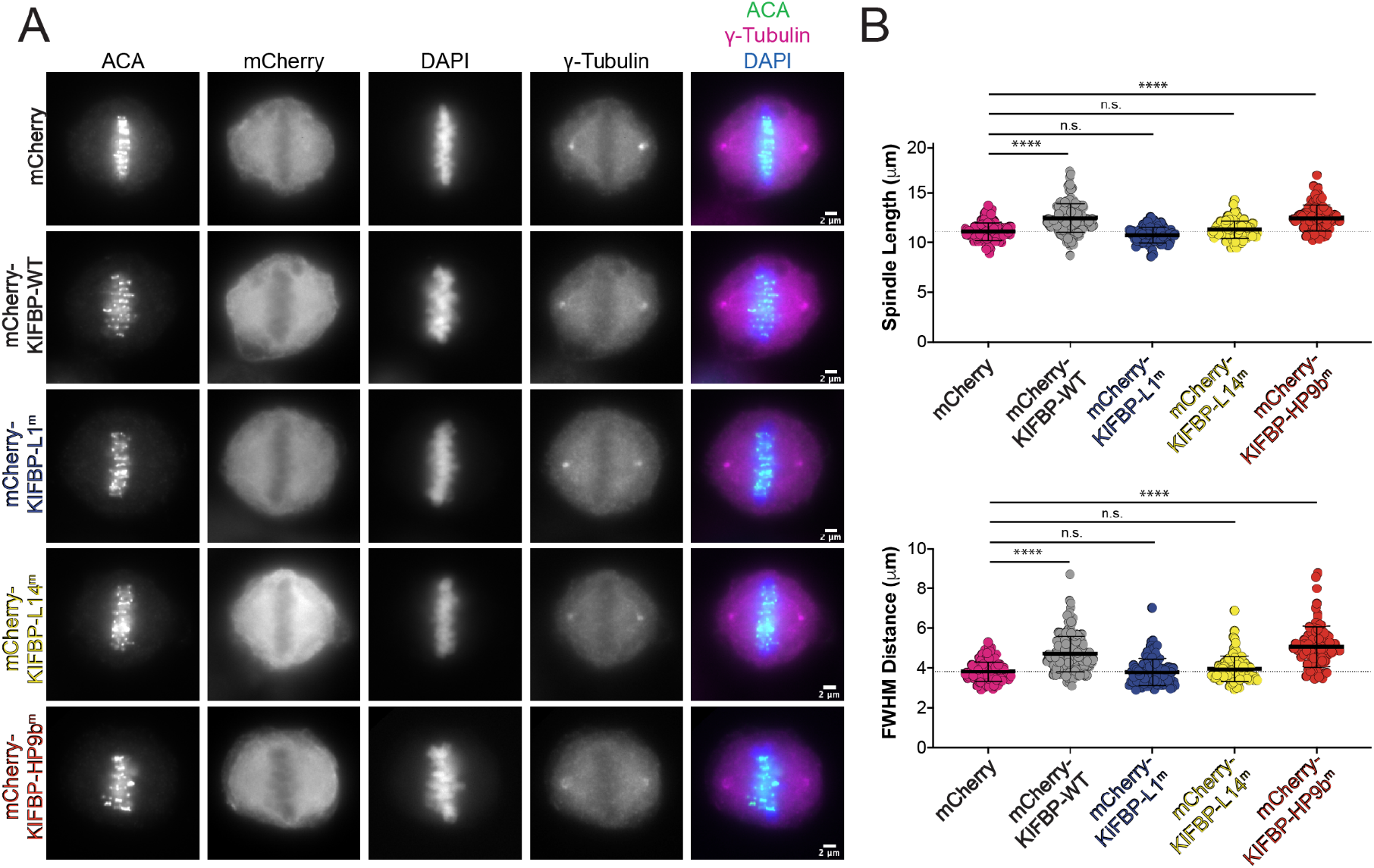
Mutations in KIFBP-L1 and KIFBP-L14 diminish KlFBP-mediated regulation of spindle length and chromosome alignment during mitosis. (A) MG132 arrested HeLa Kyoto cells overexpressing mCherry or indicated mCherry-KlFBP construct. Scale bar 2 mm. (B) Top: Graph of spindle lengths measured in cells overexpressing mCherry or indicated mCherry-KlFBP construct. Each dot represents a single cell. Mean ± standard deviation is displayed. Statistical results are shown fora one-wayANOVAwith Tukey’s Multiple Comparisons test. N.s. indicates not significant, *”* indicates adjusted p-value < 0.0001 with 95% confidence interval. See **Figure 7 - Supplement 1** for spindle length versus average mCherry expression for individual cells. Bottom: Graph of full-width at half maximum (FWHM) of centromere fluorescence distribution along the length of the spindle measured in cells overexpressing mCherry or indicated mCherry-KlFBP construct. Each dot represents a single cell. Mean ± standard deviation is displayed. Statistical results are shown for a one-way ANOVA with Tukey’s Multiple Comparisons test. N.s. indicates not significant, **** indicates adjusted p-value < 0.0001 with 95% confidence interval. See **Figure 7 - Supplement 1** for FWHM distance versus average mCherry expression for individual cells. Data were obtained from a minimum of three independent experiments. The following cell numbers were analyzed for the mCherry and mCherry-KlFBP constructs: (1) mCherry (control) = 132 cells, (2) mCherry-KlFBP-WT = 165 cells, (3) rnCherry-KIFBP-L1^m^ = 102 cells, (4) rnCherry-KIFBP-L14^m^ = 99 cells, (5) rnCherry-KIFBP-HP9b^m^ = 89 cells.

To test the mitotic effects of the KIFBP mutants, we generated mCherry-KIFBP overexpression mutant constructs mCherry-KIFBP-L1^m^, mCherry-KIFBP-L14^m^, and mCherry-KIFBP-HP9b^m^ (**Figure 6B**). Cells expressing mCherry-KIFBP-L1^m^ and mCherry-KIFBP-L14^m^ mutants displayed similar chromosome alignment and spindle lengths as cells overexpressing the mCherry control (**Figure 7A & 7B**). In contrast to mCherry-KIFBP-WT, mitotic effects of mCherry-KIFBP-L1^m^ did not scale with expression level, suggesting that mCherry-KIFBP-L1^m^ does not inhibit kinesin activity even when expressed at higher levels (**Figure 7 - Supplement 1**). Spindle length and chromosome alignment defects increased at high expression levels of mCherry-KIFBP-L14^m^, suggesting mCherry-KIFBP-L14^m^ may inhibit kinesin activity at high expression levels (**Figure 7 - Supplement 1**). In contrast to mCherry-KIFBP-L1^m^ and mCherry-KIFBP-L14^m^, mCherry-KIFBP-HP9b^m^ showed similar effects as mCherry-KIFBP-WT overexpression, decreasing chromosome alignment and increasing spindle length (**Figure 7A and 7B**). Interestingly, mitotic defects did not scale with expression level for mCherry-KIFBP-HP9b^m^, in contrast to mCherry-KIFBP-WT (**Figure 7 - Supplement 1**). Cells expressing lower levels of mCherry-KIFBP-HP9b^m^ displayed mitotic defects, suggesting that mCherry-KIFBP-HP9b^m^ may be a more potent inhibitor than mCherry-KIFBP-WT. These findings are consistent with the *in vitro* observations that KIFBP-L1^m^ and KIFBP-L14^m^ reduce KIFBP’s interaction with KIF15 and KIF18A, whereas the KIFBP-HP9b^m^ does not block interaction (**Figure 6C-E**).

To further investigate the cellular effects of the KIFBP mutations, we measured KIF18A localization in HeLa Kyoto cells overexpressing wild type KIFBP or the KIFBP mutants. KIF18A accumulates at the plus-ends of microtubules during metaphase, and we have previously shown that overexpression of mCherry-KIFBP-WT alters KIF18A spindle localization^28^. Overexpression of mCherry-KIFBP-WT abolishes KIF18A plus-end enrichment and leads to a more uniform spindle localization (**Figure 8A**), consistent with previous observations^28^. Line-scan analysis confirmed the loss of KIF18A from microtubule plus-end enrichment along individual kinetochore microtubules (**Figure 8B**).

**Figure 8.**
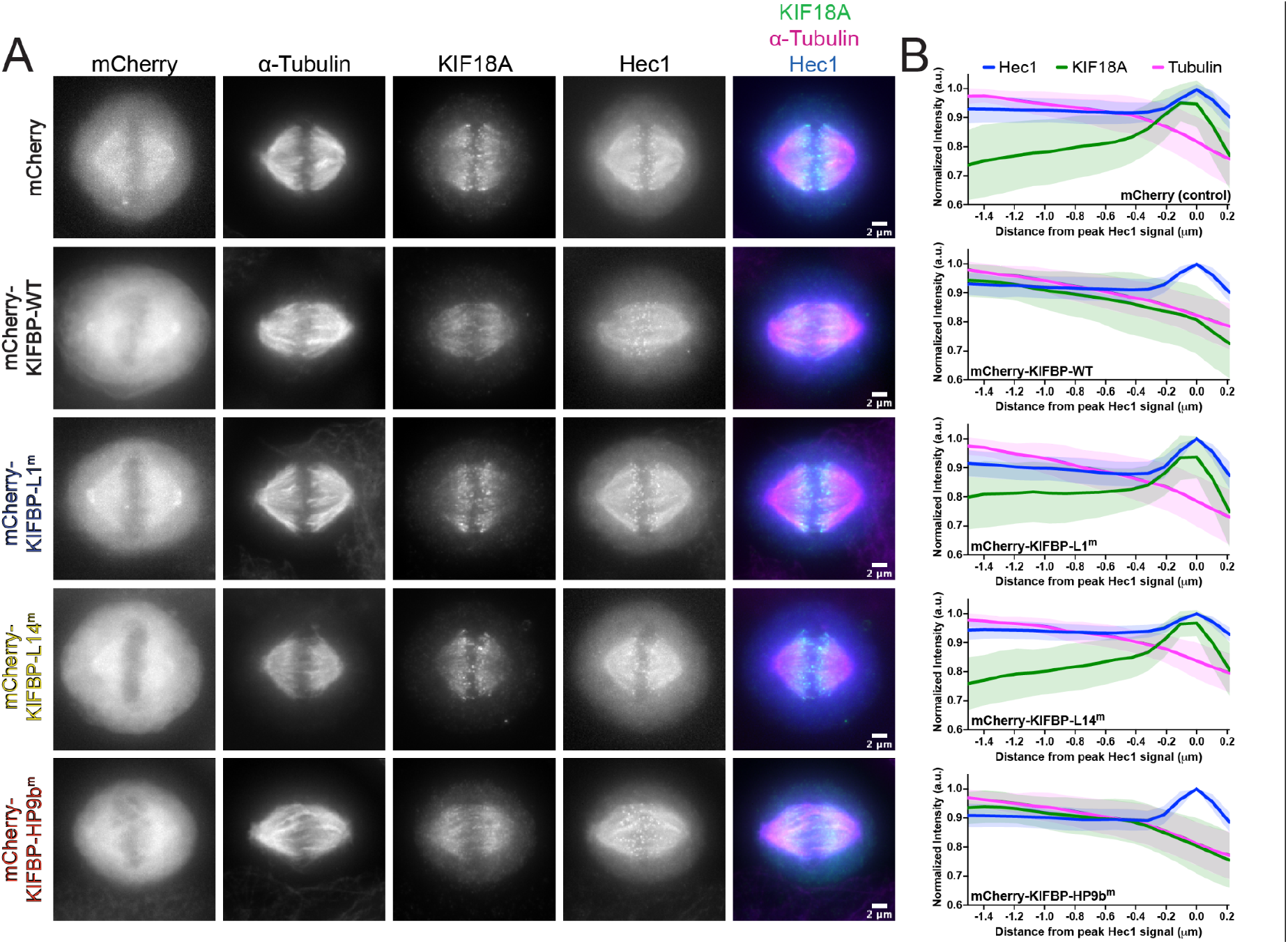
Mutations in KIFBP-L1 and KIFBP-L14 disrupt KlFBP-regulation of KIF18A localization. (A) KIF18A localization in MG132 arrested HeLa Kyoto cells overexpressing mCherry or indicated mCherry-KIFBP construct. Hed is used as a marker for the kinetochore. Scale bar 2 mm. (B) Line scan analyses of KIF18A distribution along kinetochore microtubules. Fluorescence values were normalized, aligned by peak Hed intensity, and averaged across multiple line scans. Hed, blue; KIF18A, green; Tubulin, magenta. Solid lines indicate the means, shaded areas indicate standard deviation. A.U. indicates arbitrary units. The following cell numbers and line scans were analyzed for the mCherry and mCherry-KIFBP constructs: (1) mCherry (control) = 40 cells (64 lines), (2) mCherry-KIFBP-WT = 34 cells (64 lines), (3) rnCherry-KIFBP-L1^m^ = 34 cells (64 lines), (4) rnCherry-KIFBP-L14^m^ = 32 cells (68 lines), (5) rnCherry-KIFBP-HP9b^m^ = 33 cells (63 lines).

We predicted that mutations that abolished KIFBP interaction with kinesins *in vitro* would not disrupt KIF18A localization. Overexpression of mCherry-KIFBP-L1^m^ and mCherry-KIFBP-L14^m^ did not disrupt KIF18A plus-end enrichment on kinetochore microtubules (**Figure 8A & B**). In contrast, overexpression of mCherry-KIFBP-HP9b^m^ showed similar effects to overexpression of mCherry-KIFBP-WT (**Figure 8A & B**). This is especially interesting considering that KIFBP-L14^m^ binds KIF18A *in vitro*, suggesting that the interaction is not necessarily equivalent to inhibition. Even a 10% reduction in binding may be sufficient to impair regulation by KIFBP beyond a cellular threshold. Taken together, these results support the conclusion that KIFBP Loop-1 and Loop-14 are critical regions for kinesin interaction and indicate that both regions are necessary for KIFBP to limit KIF15 and KIF18A activity during mitosis.

### KIFBP-binding kinesins adopt conformations that are distinct from non-KIFBP binding kinesins

To gain insight into how KIFBP inhibits particular kinesins, we performed a series of molecular dynamics (MD) simulations using a number of kinesin family members. We selected two members that bind KIFBP (KIF15 and KIF18A) as well as two that show little or no interaction with KIFBP (KIF5C and KIF11)^27^. Using only the motor domains, we performed 500+ nanosecond simulations of all-atom MD for each protein in the unbound, ADP state (see Methods).

To analyze the dynamics and conformational spaces explored by these proteins, we compared MD trajectories between KIFBP-binding motors (KIF15 and KIF18A) with motors that do not bind KIFBP (KIF5C and KIF11). We performed principal component analysis (PCA) using the KIF15 simulation as the reference for the other proteins to reduce dimensionality. To minimize noise in the PCA analysis that comes from fluctuations in unstructured regions of the protein, we only analyzed amino acids that were in stable secondary structure elements (ɑ-helices of β-strands) at least 80% of the time. Finally, we removed ɑ4 from the PCA analysis since this helix remained stably bound to the motors in the simulations, but is extended in the cryo-EM structure and would thus skew the PCA results.

Comparison of PCA analysis for these kinesin motors shows that 1) all four proteins have similar ranges of motion and explore comparable regions of conformational space and 2) there are specific motions that are only present in KIFBP-binding kinesins (KIF15 and KIF18A) (**Figure 9A**). Close inspection of the amino acids that contribute to PC1 (*i*.*e*., the largest motions of KIF15) revealed that splaying of both the C-terminal end of ɑ3 and the N-terminal end of ɑ6 are the dominant contributors to this mode. These structural differences are consistent with the KIFBP:KIF15 cryo-EM structure (**Figure 2D**) where movements of these helices are a defining feature of the bound complex. Thus, MD simulations indicate that only KIF15 and KIF18A are able to reach the conformation found in the bound complex (**Figure 9A**).

**Figure 9.**
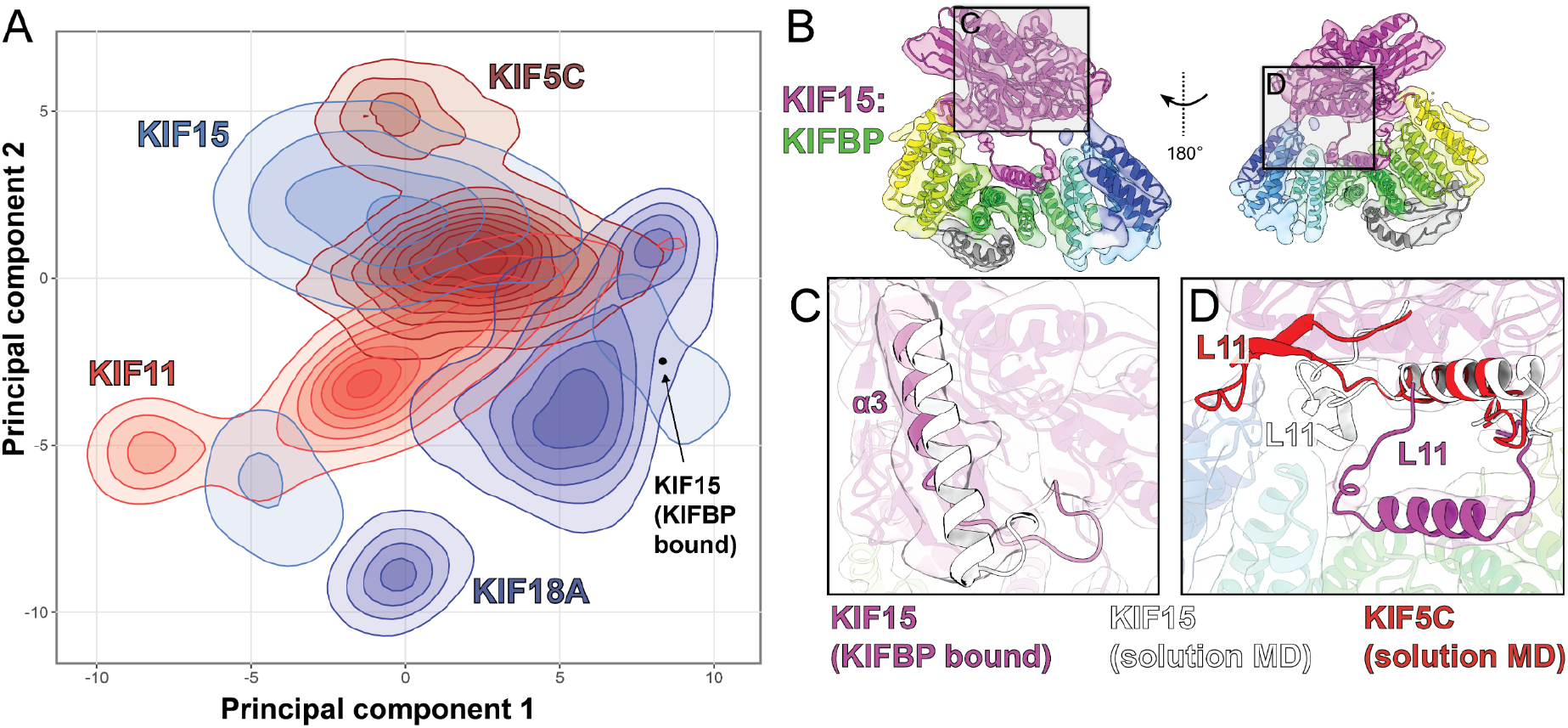
Molecular dynamics reveals specific conformations adopted by KIF15 and KIF18A that may promote KIFBP-binding. (A) Principal Component Analysis of the MD simulations for all four kinesin motors (KIF15, KIF18A, KIF5C, and KIF11). The two KIFBP binders (KIF15 and KIF18A in shades of blue) can reach the conformation of the cryo-EM complex, but the non-binders (KIF5C and KIF11 in shades of red) do not. (B) Overview of KIF15:KIFBP structure. (C) Comparison of KIF15-a3 between MD simulations and cryo-EM. (D) Differences in Loop-11 between KIF15 and KIF5C. For KIF15, L11 fits within the KIFBP pocket while the extended L11 structure of KIF5C clashes with KIFBP.

The MD KIF15 solution structure differs in specific structural regions with KIFBP-bound KIF15. First, the KIF15 solution structure shows that KIF15-ɑ4 remains closely bound to the motor body throughout each simulation. This would suggest that the release and translocation of KIF15-ɑ4 would only occur upon interaction with KIFBP, or on a much longer time scale than that sampled during the simulation. Second, even though we saw that the movement of KIF15-ɑ3 was a key feature from the PCA analysis, we see in the cryo-EM structure that the C-terminal end of KIF15-ɑ3 loses its structure over the final 8-9 residues, forming an extended loop with a short antiparallel β-sheet (**Figure 9B & C**). This part of KIF15 makes contact with KIFBP and it appears that this structural change would allow better contact between KIF15 and KIFBP.

Lastly, even though ɑ4 has a nearly identical conformation for all kinesins, we see that L11 is much more dynamic and its conformation is kinesin-dependent. For KIF5C, L11 tends to be extended away from ɑ4, and when superimposed on the cryo-EM complex, we see that L11 has a steric clash with KIFBP (**Figure 9B & D**). Conversely, both KIF15 and KIF18A adopt more compact L11 structures and these fit well within the cavity of the KIFBP structure (KIF15 structure shown in **Figure 9D**). Taken together, this suggests that the overall conformation of the motor head as well as the dynamics or conformation of more flexible parts of each protein may act in concert to determine which kinesins will bind to KIFBP and which will not.

## DISCUSSION

Our work presents a new mode of kinesin motor protein regulation *via* a multivalent interaction between KIFBP and the kinesin motor domain. Using a combination of cryo-EM, XL-MS, MD simulations, biochemical assays, and cell biology, we describe a model in which KIFBP stabilizes the microtubule-binding ɑ4 helix^5,40–42^ away from the kinesin motor domain to ensure complete inhibition of kinesin microtubule-binding. Interestingly, KIFBP does not mimic the negatively-charged microtubule surface^43^ to engage kinesin motors. Instead, KIFBP utilizes a hydrophobic cleft to hold ɑ4 and Loop-11 in a conformation that is incompatible with microtubule-binding, while simultaneously engaging and sterically inhibiting the kinesin microtubule-binding interface (**Figure 3**).

Determining the binding mechanism of KIFBP relied on cryo-EM structural data for relatively small macromolecular samples. We determined a near-atomic structure of KIFBP corresponding to ∼40kDa out of 72 kDa, using this reconstruction for *de novo* model building. This places the KIFBP reconstruction and model among the smallest molecular weight macromolecules to be built *de novo* by cryo-EM. The relatively small size of KIFBP may be the driving factor limiting the overall resolution of KIFBP alone to ∼3.8Å instead of obtaining higher-resolution reconstructions.

During the preparation of this work, another study utilized cryo-EM and cell biology assays to propose a mechanism of KIFBP-mediated kinesin inhibition^38^. In this study, Atherton et al. determined lower resolution structures of KIFBP (4.8Å) and KIFBP-KIF15 (6.9Å) to arrive at a similar model of kinesin motor inhibition, where KIFBP stabilizes the ɑ4 helix away from KIF15. In our work, our higher-resolution structures of KIFBP alone (3.8Å) and KIFBP-KIF15 (4.8Å) allowed us to 1) build a high-confidence atomic model for KIFBP and 2) map conformational changes in the KIF15 motor when bound to KIFBP. Importantly, we showed that KIFBP utilizes a similar mode of inhibition for KIF18A, indicating that our proposed model is likely a general mode of kinesin inhibition. Finally, we used our structures alongside XL-MS to map the interaction of KIFBP with kinesin motors and showed that blocking the interaction between KIFBP with KIF15 and KIF18A *via* mutagenesis minimizes its ability to regulate motor activity in the physiologically-relevant context of mitosis (**Figures 7 & 8**).

### Kinesin motor recognition by KIFBP

Our cryo-EM reconstruction of KIFBP reveals that KIBP contains a 9-TPR array, which folds into a solenoid with a concave kinesin-interacting surface. Unlike continuous TPR proteins like LGN^44^, KIFBP is punctuated by a centrally located helix and loop (**Figure 1**). The binding of KIFBP to kinesin motor heads is strikingly different from the interaction of many TPR proteins to their ligands^34^. Many TPR proteins bind a short sequence, e.g. HOP binds the motif MEEVD in Hsp90^45^. In contrast, our cryo-EM and XL-MS show that the interaction of KIFBP with kinesin motors is highly multivalent. First, KIFBP-L1, localized at its N-terminus, engages ɑ3 and β6 of the kinesin motor head. Second, KIFBP-L14 contacts β4-β5 of KIF15 and KIF18A with amino acid E168 in the KIF15 motor domain (or E161 in KIF18A) positioned to play a key role in this interaction. Third, ɑ4 helices of KIF15 and KIF18A become nestled into a multi-helix groove created by KIFBP-HP4a, -HP4b, and -HP5a. The binding of KIF15/KIF18A-ɑ4 to KIFBP is striking because it requires a 15 Å displacement of the helix from its resting position within the kinesin motor head.

Our structure- and XL-MS-guided mutagenesis study of KIFBP revealed that KIFBP-L1 is especially important for motor binding. Charge-neutralization of Loop-1 renders KIFBP incompetent for binding both KIF15 and KIF18A *in vitro* (**Figure 6**). Consistent with these data, the KIFBP-L1^m^ failed to produce phenotypes associated with KIFBP overexpression in cells, *i*.*e*., disruption of chromosome alignment and increased spindle length (**Figures 7 & 8**). Unlike KIFBP-WT, the KIFBP-L1^m^ mutant was also incapable of disrupting KIF18A localization.

In contrast to KIFBP-L1, the role of KIFBP-L14 in motor inhibition remains less clear. Previous work utilizing an artificial peroxisome transport assay suggested that KIFBP-L14 is important for the ability of KIFBP to inhibit the motor activities of KIF1A and KIF15^38^. Interestingly, however, mutation of KIFBP-L14 more strongly affected the ability of KIFBP to retrieve KIF1A from cell lysates than KIF15. In our experiments with purified proteins, we observed that KIFBP-L14^m^ showed reduced binding to KIF15, but barely reduced interaction with KIF18A (**Figure 6**). Surprisingly, in cells, KIFBP-L14^m^ reduced the ability of KIFBP to disrupt metaphase chromosome alignment, spindle length homeostasis and KIF18A localization (**Figures 7 & 8**). However, high levels of KIFBP-L14^m^ did produce phenotypes associated with KIFBP overexpression, suggesting that KIFBP-L14^m^ is capable of weak interactions with KIF18A and/or KIF15. Taken together, these data suggest KIFBP-L14 may be more important for binding some kinesins than others, but that binding for all three kinesins collectively tested (KIF15, KIF18A, and KIF1A) was reduced beyond some cellular threshold necessary for producing the observed phenotypes of mitotic defects or reduced peroxisome transport. More work is required to relate the ability of KIFBP to bind kinesin motors in vitro to its ability to regulate kinesins in cells.

Lastly, we observed a high density of crosslinks between KIFBP-HP9b (residues K610 and K617) and the microtubule-binding interface of KIF15 (**Figure 4**). These structural elements are not within the crosslinking range of BS3 in our cryo-EM structures (**Figures 2 & 5**), and the significance of these crosslinks is therefore not clear. Perhaps in line with this, our analysis of the charge-neutralization mutant KIFBP-HP9b^m^ revealed that this mutation had little effect on *in vitro* interaction with KIF15 or KIF18A or on the mitotic phenotypes we quantified, suggesting that electrostatic interactions with amino acids 610-617 of KIFBP are not critical for kinesin interaction. A role for the C-terminus in KIFBP-motor interactions should not be dismissed, as a recent study identified a novel nonsense KIFBP mutation in a GOSHS patient that truncates the protein at position 593^46^. It will be interesting to determine if the C-terminus of KIFBP is generally important for its interaction with all kinesins, or if it instead drives interactions with kinesins that are more clinically relevant to GOSHS.

### KIFBP remodels the kinesin motor head to displace kinesin-α4

Kinesin ɑ4 helix plays a critical role in motor-KIFBP binding. Lysine residues within KIFBP-HP4a (K205) and KIFBP-LH (K307) crosslink residues located in KIF15-L11 (K273 and K283). Surprisingly, residues in KIFBP-L1 (K26, K30, K36) crosslinked KIF15 residues K273 and K283. The significance of these crosslinks is not clear, but these data may suggest that an intermediary complex between KIFBP and kinesin motor domains, driven by the interaction of KIFBP-L1 with kinesin Loop11, may form prior to the acquisition of the final bound state.

One outstanding question concerns the mechanism by which KIF15/KIF18A ɑ4 undergoes long-range motion to achieve KIFBP binding. The simplest possibility is that ɑ4 is positionally unstable. If sufficiently compliant, the adjacent loops, *i*.*e*., Loops11 and 12, may allow ɑ4 excursions that eventually result in “capture” of ɑ4 by KIFBP. Long-range motions of ɑ4 are not without precedent. For example, Wang *et al*. (2016) observed by X-ray crystallography that ɑ4 of KIF19 is positioned much farther from the motor head than is typical^47^. Our MD work, however, suggests that ɑ4 remains closely associated with the motor head (data not shown), leading us to speculate that the binding of a motor head by KIFBP results in allosteric changes in the structure of both proteins, inducing motions of ɑ4 that predispose it to KIFBP binding. Comparison of KIF15 in apo *versus* bound states supports this possibility (**Figures 2D-E**). When bound to KIFBP, KIF15 showed a shift of several alpha helices (α1, α3, α6) away from the core of the motor as well as large movements of several beta-strand pairs. The movement of these structural elements causes the motor to assume a more open conformation.

In summary, our work establishes a structural mechanism by which KIFBP inactivates the microtubule-binding activity of mitotic kinesins KIF15 and KIF18A. Unlike common TPR tandem proteins, KIFBP uses multivalency to form a complex with the kinesin motor head. Multivalency may explain why it has not been possible to identify a consensus sequence for kinesin motors that bind KIFBP *versus* those that do not^38^. Our MD simulations and PCA analysis also indicate that motor-specific steric clashes may serve as a mechanism that prevents certain motors from binding KIFBP. Specifically, we observed that L11 of KIF5C would sterically clash with KIFBP-HP2, whereas L11 of KIF15 and KIF18A fits within the cavity between HP3 & HP4 of KIFBP. Further work is required to test the generality of this idea. An additional area for future work is to reveal the mechanism by which KIFBP dissociates from a kinesin motor. The multivalency with which KIFBP interacts with a kinesin motor, in particular the interaction of HP4a/b-HP5 with ɑ4, suggests that a motor will not readily disengage from KIFBP. Motor recycling may require active regulation, e.g., phosphorylation, as proposed in earlier work^27^.

## ACKNOWLEDGEMENTS

We thank members of the Michigan Cytoskeleton Supergroup for helpful discussions and critical feedback. The authors thank Dr. Venkatesha Basrur of the Proteomics Resource Facility at the University of Michigan Medical School for providing technical expertise in mass spectrometry. Research reported in this publication was supported by the University of Michigan Cryo-EM Facility (U-M Cryo-EM). U-M Cryo-EM is grateful for support from the U-M Life Sciences Institute and the U-M Biosciences Initiative. This work was supported by start-up funds from the University of Michigan (R.O., M.A.C.); NIH grants GM094231 (A.I.N.), GM136822 (D.S.), GM121491 (J.S.), and GM086610 (R.O.). L.J. is currently an IRACDA fellow supported by the NIH (K12 GM111725). The research reported in this publication was supported by the NIH under award No. S10OD020011.

## AUTHOR CONTRIBUTIONS

R.O., A.L.S., J.S., and M.A.C. conceived the project. A.L.S. performed protein purifications; protein crosslinking; site-directed mutagenesis of KIFBP; and protein-protein interaction assays in vitro. Z.T. performed cryo-electron microscopy of KIFBP, KIF15, and KIF18A, data collection, and analysis. Z.T. and M.A.C. analyzed cryo-EM data, built the structure of KIFBP from its primary sequence de novo, and analyzed protein conformational changes. S.E.H. and A.L.S. analyzed crosslinking mass spectrometry data. K.L.S. created KIFBP mutant constructs and analyzed the effects of their overexpression in cells. L.J. and D.S. performed and analyzed the molecular dynamics simulations. A.L.S., K.L.S., D.S., J.S., R.O., and M.A.C. wrote the paper with input from all authors.

## MATERIALS & METHODS

### Plasmid construction

The following plasmids that were used in this study were previously described elsewhere: GST-KIFBP^28^, mCherry and mCherry-KIFBP expression plasmids^28^.

His_6_-KIF15-N375 was created through isothermal assembly where the first 375 amino acids of the KIF15 open reading frame were amplified from pEGFP-C1-Kif15-FL^48^ and inserted into the pET15b vector. Correct insertion was confirmed by sequencing.

His_6_-KIF18A-N363 was created through isothermal assembly where a gBlock gene fragment of the first 363 amino acids of KIF18A codon-optimized for expression in E. coli (IDT) was inserted into the pET15b vector. Correct insertion was confirmed by sequencing.

GST-KIFBP-L1^m^ was created by site-directed mutagenesis of GST-KIFBP, replacing amino acids 21-40 with the altered amino acid sequence described in **Figure 6B**. Similarly, GST-KIFBP-HP9b^m^ was created by site-directed mutagenesis of GST-KIFBP replacing amino acids 610-617 with the altered amino acids sequence described in **Figure 6B**. Mutagenesis was confirmed by sequencing of the open reading frame. mCherry-KIFBP-L1^m^ and mCherry-KIFBP-HP9b^m^ were created in the same manner by site-directed mutagenesis of the mCherry-KIFBP wild-type plasmid using the primers described in the Key Resources Table. PCR products were circularized using the commercially available KLD Enzyme Mix (New England Biolabs). The resulting plasmids were confirmed by sequencing.

To create the mCherry-KIFBP-L14^m^ plasmid, a GeneStrand containing KIFBP base pairs 1092-1588 with the L14^m^ mutations was synthesized (Eurofins) (Sequence provided in Key Resources Table). This gene fragment was then inserted into the mCherry-KIFBP expression vector by isothermal assembly using the commercially available Gibson Assembly Master Mix (New England Biolabs) after PCR amplification of the mCherry-KIFBP expression vector with the primers described in the Key Resources Table. The resulting clones were confirmed by sequencing. GST-KIFBP-L14^m^ was then created by isothermal assembly wherein the full KIFBP sequence containing the mutated residues was inserted into the pGEX-6P-1 vector. Correct insertion was confirmed via sequencing.

### Protein expression and purification

Expression of GST-KIFBP, GST-KIFBP-L1^m^, GST-KIFBP-L14^m^, and GST-KIFBP-HP9b^m^ were induced in BL21-DE3 cells with 0.4 M IPTG overnight at 16° C. Cells were pelleted and resuspended in lysis buffer (1X PBS, 0.5 mM NaCl, 5 mM β-mercaptoethanol, 1% NP-40, and protease inhibitors [1 mM PMSF, 1 mM benzamidine, and 10 ug/mL LPC]), after which they were incubated with 1 mg/mL lysozyme for 30 minutes on ice followed by sonication. The lysate was clarified by centrifugation for 30 min at 35,000 rpm at 4°C in a Type 45 Ti rotor (Beckman). Cleared lysate was incubated with 2 mL glutathione-sepharose (Fisher Scientific) for 1 hr and washed with 50 mL (25 CV) wash buffer (1X PBS, 0.5 M NaCl, 5 mM β-mercaptoethanol). Resin was incubated with 200 uL Precission Protease (Cytiva) in 2 mL cleavage buffer (50 mM Tris-HCl, pH 7.0, 150 mM NaCl, 1 mM EDTA, 1 mM DTT) for 4 hr at 4° C to cleave the GST tag. Protein was eluted with 50 mM Tris-HCl, pH 8.0, and peak fractions were combined and clarified by centrifugation for 5 min at 20,000 rpm at 4°C, after which they were subjected to size exclusion chromatography on a Superdex 200 column (GE Healthcare) equilibrated in 10 mM K-HEPES, pH 7.7, 50 mM KCl, and 1 mM DTT. Protein concentration of fractions after gel filtration were estimated using a Bradford assay, after which peak fractions were combined, concentrated to >1 mg/mL using Amicon 10 kDa centrifugal filter units (Millipore), and either used immediately for cryo-EM or flash frozen and stored at -80°C.

Expression of His_6_-KIF15-N375 and His_6_-KIF18A-N363 was induced in BL21-DE3 cells with 0.4 M IPTG overnight at 16° C. Cells were pelleted and resuspended in lysis buffer (1X PNI [50 mM sodium phosphate, 500 mM NaCl, 20 mM imidazole], 1% NP-40, 1 mM MgATP, and protease inhibitors [1 mM PMSF, 1 mM benzamidine, and 10 ug/mL LPC]), after which they were incubated with 1 mg/mL lysozyme for 30 minutes on ice followed by sonication. The lysate was clarified by centrifugation for 30 min at 35,000 rpm at 4°C in a Type 45 Ti rotor (Beckman). Cleared lysate was incubated with 2 mL Ni-NTA agarose (Qiagen) for 1 hr and washed with 50 mL wash buffer (1X PNI, 100 µM MgATP, 5 mM β-mercaptoethanol). Protein was eluted with elution buffer (1X PNI, 100 µM MgATP, 5 mM β-mercaptoethanol, 200 mM imidazole) and peak fractions were combined and clarified by centrifugation for 5 min at 20,000 rpm at 4°C, after which they were subjected to size exclusion chromatography on a Superdex 200 column equilibrated in gel filtration buffer (10 mM K-HEPES, pH 7.7, 50 mM KCl, 1 mM DTT, and 0.2 mM MgATP). Protein concentration of fractions after gel filtration were estimated using a Bradford assay, after which peak fractions were combined and concentrated to >1 mg/mL using Amicon 10 kDa centrifugal filter units (Millipore). Prior to cryo-EM grid preparation, the protein was then mixed with equimolar KIFBP and subjected to size exclusion chromatography a second time on a Superose 6 column (GE Healthcare) equilibrated with gel filtration buffer. Peak fractions were analyzed by SDS-PAGE and stained with Coomassie blue. Fraction(s) containing only the two proteins of interest were then combined, concentrated to >1 mg/mL, and used immediately for cryo-EM.

### Cryo-EM grid preparation and data collection

For cryo-EM grid preparation, after size exclusion chromatography, KIFBP was concentrated to 4mg/ml whereas KIFBP:KIF15 and KIFBP:KIF18A complexes were each concentrated to 1mg/ml. Aliquots of 4ul were applied on glow-discharged UltraAuFoil R(1.2/1.3) 300 mesh gold grids (Electron Microscopy Sciences). The grids were then blotted with filter paper and plunge-frozen into liquid ethane cooled by liquid nitrogen using a Vitrobot Mark IV (Thermo Fisher Scientific) set to 4°C, 100% humidity, 1.5s blot, and a force of 20.

For KIFBP and KIFBP:KIF15 samples, datasets were collected using Leginon^49^ on a ThermoFisher Glacios transmission electron microscope operating at 200keV equipped with a Gatan K2 Summit direct electron detector (Gatan Inc.) in counting mode. For KIFBP, a total of 11,086 micrographs were collected through 3 data collection sessions with total doses of 58-68e^-^/Å ^2^ during exposure time of 7-9s, dose fractionated into 35-45 movie frames at defocus ranges of 1-2μm. The magnification used here is 45000x, resulting in a physical pixel size of 0.98 Å per pixel. For KIFBP:KIF15, 4 data collection sessions were performed with 45000x magnification and a physical pixel size of 0.98 Å per pixel. A total number of 6,184 micrographs were collected with a total dose of 60 e^-^/Å ^2^ during 6-7s exposure time, dose fractionated into 30-35 movie frames.

Data collection for the KIFBP:KIF18A sample was automatically collected using Leginon^49^ on an FEI Talos Arctica transmission electron microscope operating at 200keV equipped with a Gatan K2 Summit direct electron detector in counting mode. Three datasets were collected, resulting in a total 4,669 micrographs in a physical pixel size of 0.91 Å per pixel. The total dose ranges from 52-62 e^-^/Å ^2^ in a 7-8s exposure time with dose fractionated into 35-40 movie frames.

### Cryo-EM data processing

The data processing diagram for KIFBP is shown in **Figure 1 - Supplement 2 and 3**. Movie alignment, CTF parameter estimation, and particle picking were performed using Warp^50^. The resulting particles were imported into cryoSPARC^51^ and underwent iterative 2D classification to remove bad classes. We initially analyzed sample heterogeneity in ‘dataset3’ using *ab-initio* reconstruction with 3 classes. Particles from the one good class were then subjected to another round *ab-initio* reconstruction with 2 classes, resulting in two very similar classes. After careful examination, we believe that the first class is the full-length KIFBP map, while the other is the KIFBP lack of the C-terminal helices, explaining why some class averages are missing the C-terminal helices pairs of the KIFBP.

To resolve the structure of the full KIFBP, we deliberately selected the class averages from ‘dataset1’ and ‘dataset2’ that resemble the full KIFBP molecule using cryoSPARC^51^. These particles were then combined with the particles from the full-length KIFBP class in ‘dataset3’ and subjected to *ab-initio* reconstruction with 2 classes. Particles from the good classes were selected for non-uniform refinement^52^ to obtain a 4.7Å resolution map. The map quality was improved to 4.6Å using local refinement with a static mask. The resulting map was manually sharpened for the visual inspection purpose using a B-factor of -50Å ^2^.

Next, we focused on analyzing the N-terminus of KIFBP lacking the C-terminal helices to try to improve the resolution. Particles from ‘dataset2’ were chosen since CTF fit resolution in ‘dataset2’ was the best among all three datasets. 913,455 particles from Warp^50^ were imported into cryoSPARC^51^. We then performed iterative heterogeneous refinement with one good class from the KIFBP lacking the C-terminal helices and two bad classes from failed *ab-initio* reconstruction jobs. 301,491 particles corresponding to KIFBP were enriched after extensive heterogeneous refinement and 2D classification. These particles were then subjected to homogeneous refinement and local refinement, resulting in a 3.8Å resolution map. However, the quality of the map was not satisfactory. To improve the map quality, the particles were exported into RELION-3.1^53^ and underwent one round of 3D auto-refinement to obtain a reconstruction at 4.2Å. Subsequently, two rounds of CTF refinement were performed to correct for beam tilt^54^. 3D auto-refinement with refined beam-tilt yielded an estimated resolution of 4.1Å. We then exported micrograph motion trajectories from Warp and performed Bayesian polishing^55^ to optimize per-particle motion tracks. 3D auto-refinement from the polished particles resulted in a 3.8Å map. Following this step, one round of 3D classification with 6 classes was performed to further remove heterogeneity. 5 classes corresponding to KIFBP lacking the C-terminus were selected and underwent another round of refinement, yielding a 3.8Å resolution map. The particles were then re-extracted and re-centered. The following 3D auto-refinement yielded a 4Å map. Bayesian polishing was performed on this particle stack, resulting in an improved map quality. 3D classification without alignment using T=12 resulted in one high-resolution class. 3D auto-refinement using the 128,190 particles from this high-resolution class gave a 3.8Å map with improved map quality.

For KIFBP:KIF15, all the data processing steps were performed in cryoSPARC^51^, as presented in **Figure 2 - Supplement 3**. To generate an initial map for the KIFBP:KIF15 complex, 1,007 movies from dataset1 were imported into cryoSAPRC^51^. Patch motion correction and patch CTF estimation were used to correct beam-induced motion and estimate CTF parameters. 235,721 particles were automatically picked using the Topaz general model^56^. These particles were then subjected to iterative 2D classification to remove bad classes. The resulting 32,262 particles were used for *ab-initio* reconstruction with one class and also retraining Topaz. The model from the ab-initio reconstruction was refined to ∼7Å and used as a template for the heterogeneous refinement in the following steps.

Movies from dataset1, 2, 3, and 4 were imported into cryoSPARC^51^ and processed separately at the beginning steps. Movies were aligned using patch motion correction with dose weighting. CTF parameters were estimated with patch CTF estimation. Micrographs with CTF fit resolution below 5Å were selected and subjected to particle picking using a restrained Topaz model. The picked particles underwent one round 2D classification to remove obvious junk. We then performed iterative heterogeneous refinement with one good class from the initial template and two bad classes from the early-terminated *ab-initio* reconstruction jobs to enrich particles corresponding to KIFBP:KIF15 complex. The resulting particles were further cleaned by 2D classification and *ab-initio* reconstruction with multi-classes. Particles from the individual dataset were then re-extracted, re-centered and combined, resulting in 189,984 particles. These particles were subsequently classified into three classes using *ab-initio* reconstruction. Two classes showing KIFBP and KIF15 density were merged and further classified into two classes with *ab-initio* reconstruction. One class with the better KIF15 motor domain density was selected and subjected to homogenous refinement, resulting in a 4.8Å resolution map. Then local refinement with a user-defined mask was performed to improve the map quality.

For KIFBP:KIF18A, the data processing diagram is presented in **Figure 5 - Supplement 3**. A total 4,669 micrographs were collected through three datasets. For each dataset, motion correction, CTF estimation and particle picking was performed in Warp^50^, resulting in 71,529 particles (dataset1), 94,716 particles (dataset2) and 638,186 particles (dataset3). These particles were imported into cryoSPARC^51^ and underwent interactive 2D classification to remove junks. The remaining 156,159 particles were used for *ab-initio* reconstruction into four classes in cryoSPARC. 54,801 particles from the class with clear KIF18A density were selected for homogeneous refinement to obtain a 5 Å resolution structure of the KIFBP:KIF18A complex.

The quality of the map was further improved by local refinement in cryoSPARC with a user-defined mask.

### Model building

To construct an atomic model of KIFBP, first, we began by *de novo* building of the 3.8Å KIFBP reconstruction using Coot^57^ on the RELION^58^ post-processed reconstruction. To guide model building, we used density modification with DeepEMhancer^59^ that was run on the COSMIC^2^ science gateway^60^ to help interpret the cryo-EM density. From this process, we built amino acids 5-403 of KIFBP and the RELION post-processed map was used for model refinement and validation using Phenix^61^. After building this high-resolution part of our reconstruction, we built poly-alanine models for the C-terminal helices KIFBP-HP6b, -HP7, -HP8, and HP9 using Coot^59^. The manual build model was then subjected to real-space refinement in Phenix^61^.

Due to the moderate resolution (4.8Å) of KIF15:KIFBP, we built the model of KIF15:KIFBP using a combination of Rosetta-CM^35^, Rosetta-Relax, and manual building in Coot^57^. For the KIFBP model, we manually docked the KIFBP model into the density using Chimera^62^ after which we fit the model into the density Rosetta-Relax. To fit KIF15 into the density, we manually docked KIF15 (PDB: 4BN2)^36^ into the cryo-EM density, removing KIF15-L11,-α4, and -L12 from the model. With this docking, we then ran Rosetta-CM^35^, using atomic models 1V8K (Chain A)^63^, 2OWM (Chain B)^64^, 3U06 (Chain A)^65^, 4BN2 (Chain C)^36^, 5GSZ (Chain A)^66^, 5MIO (Chain C)^67^, 5MLV (Chain D)^68^, 5MM4 (Chain K)^69^, 5MM7 (Chain K)^69^, 6B0I (Chain K)^70^ as the library of fragments for rebuilding. After running Rosetta-CM to calculate 5000 models, we used the lowest scoring model for the final step of Rosetta-Relax. To build KIF15-L11,-α4, and -L12, we built a polyalanine model manually using Coot^57^.

For the KIF18A:KIFBP model, we used Rosetta-Relax to fit the KIFBP model into the density. The KIF18A motor, we manually docked the crystal structure of KIF18A (PDB: 3LRE)^71^ into the density with the exception of KIF18A-L11,-α4, and -L12. To build KIF18A-L11,-α4, and -L12, we built a polyalanine model manually using Coot^57^.

The efficiency (cryoEF)^72^ for each reconstruction was calculated on the COSMIC^2^ science gateway^60^. Figures were prepared using Chimera^62^ and ChimeraX^73^.

### Crosslinking mass spectrometry

His_6_-KIF15-N375 and KIFBP were purified as described above. An equimolar solution of both proteins was prepared in a crosslinking buffer (40 mM HEPES pH 7.4) where the total protein concentration was 10 uM and the amount of each protein was at least 20 ug. A 50 mM solution of the 11 Å lysine-targeting crosslinker BS3 was prepared in water and added to the reaction in a 100-molar excess. The reaction proceeded for 30 min while rotating at 4° C, after which it was quenched with Tris-HCl pH 7.5 at a final concentration of 50 mM. As an un-crosslinked control, a separate reaction was prepared and quenched the same way but no crosslinker was added.

The crosslinking reactions were resuspended in 50 uL of 0.1M ammonium bicarbonate buffer (pH∼8). Cysteines were reduced by adding 50 uL of 10 mM DTT and incubating at 45°C for 30 min. Samples were cooled to room temperature and alkylation of cysteines was achieved by incubating with 65 mM 2-Chloroacetamide, under darkness, for 30 min at room temperature. Overnight digestion with 1:50 enzyme:substrate modified trypsin was carried out at 37°C with constant shaking in a Thermomixer. Digestion was stopped by acidification and peptides were desalted using SepPak C18 cartridges using the manufacturer’s protocol (Waters). Samples were completely dried via vacufuge. Resulting peptides were dissolved in 9 uL of 0.1% formic acid/2% acetonitrile solution, and 2 uL of the peptide solution were resolved on a nano-capillary reverse phase column (Acclaim PepMap C18, 2 micron, 50 cm, ThermoScientific) using a 0.1% formic acid/2% acetonitrile (Buffer A) and 0.1% formic acid/95% acetonitrile (Buffer B) gradient at 300 nl/min over a period of 180 min (2-25% buffer B in 110 min, 25-40% in 20 min, 40-90% in 5 min followed by holding at 90% buffer B for 10 min and equilibration with Buffer A for 30 min). The eluent was directly introduced into a Q exactive HF mass spectrometer (Thermo Scientific, San Jose CA) using an EasySpray source. MS1 scans were acquired at 60K resolution (AGC target=3×106; max IT=50 ms). Data-dependent collision-induced dissociation MS/MS spectra were acquired using the Top speed method (3 seconds) following each MS1 scan (NCE ∼28%; 15K resolution; AGC target 1×105; max IT 45 ms).

pLink v2.3.9 was used to perform database searching against a FASTA protein sequence file containing full-length KIF15, KIFBP, and 292 common contaminant proteins. Raw data files were searched with BS3 as the crosslinker, Trypsin_P allowing up to 3 missed cleavages, peptide mass between 500 and 6000, peptide length between 5 and 60, precursor and fragment tolerances set to 20 ppm, fixed carbamidomethyl cysteine, variable methionine oxidation, 10 ppm filter tolerance, and separate 5% PSM FDR. pLabel v2.4.1 was used to visualize crosslinked MS/MS spectra. Of the resulting FDR-filtered list of crosslinked peptides, we filtered out all intra-KIFBP and intra-KIF15 crosslinks, as well as crosslinks with contaminant proteins and crosslinks with an e-value >0.05.

### *In vitro* pull-down assays

100 μL of Ni-NTA Agarose resin (Qiagen) per condition was equilibrated with binding buffer (10 mM HEPES pH 7.4, 50 mM KCl, 10 mM imidazole) and incubated with 250 μg of either His_6_-KIF15-N375 or His_6_-KIF18A-N363 for 30 min at 4°C. The kinesin-bound resin was then washed 3 times in batch with 10 CV of binding buffer, split into 20 uL aliquots, and incubated with 20 μg of either KIFBP-WT, KIFBP-L1^m^, KIFBP-L14^m^, KIFBP-HP9b^m^, or GST in a total volume of 200 μL for 30 min at 4°C. To control for non-specific binding to the resin, 20 μL of resin was incubated with either 20 μg of WT-KIFBP or GST with no previous kinesin-incubation step. After incubation with GST or KIFBP proteins, the resin was pelleted and the supernatant was removed and saved for analysis. The resin was then washed 5 times with 10 CV of wash buffer (binding buffer with 0.05% Tween-20). After the final wash, each resin sample was resuspended in 80 μL of binding buffer, and samples were taken for analysis. 5% of each supernatant and pellet sample was boiled in 5X SDS-sample dye and loaded onto a 10% SDS-PAGE gel. Gels were stained with Coomassie-blue and quantifications of band intensities were done with ImageJ. Images for publication were enhanced with ImageJ, while quantification was done from raw images.

### Cell culture and transfections

HeLa Kyoto cells were cultured at 37°C with 5% CO_2_ in MEM-α medium (Gibco) containing 10% Fetal Bovine Serum (FBS) (Gibco). For plasmid transfections in a 24-well plate format, ∼75,000 cells in 500 μl MEM-α medium were seeded onto acid-washed glass coverslips and subsequently transfected with 375 ng mCherry alone or mCherry-KIFBP plasmid DNA (containing wild type KIFBP sequence or indicated KIFBP mutant). Cells were treated with mCherry and indicated mCherry-KIFBP plasmids that were preincubated for 10 minutes in 50 μl Opti-MEM (Gibco) and 1 μl Lipofectamine LTX reagent (Invitrogen). Plasmid transfections were incubated for 24 hours before fixation for immunofluorescence.

### Cell fixation and immunofluorescence

For metaphase observations of spindle length and chromosome alignment, cells expressing mCherry and mCherry-KIFBP (wild type or indicated mutants) were treated with 20 μM MG132 (Selleck Chemicals) for 2 hours before fixation. Cells were fixed on coverslips in -20°C methanol (Fisher Scientific) with 1% paraformaldehyde (Electron Microscopy Sciences) for 10 minutes on ice. Coverslips were then washed three times for 5 minutes each in Tris-Buffered Saline (TBS; 150 mM NaCl, 50 mM Tris base, pH 7.4). Coverslips were blocked for 1 hour at room temperature in 20% goat serum in antibody dilution buffer (Abdil: TBS pH 7.4, 1% Bovine Serum Albumin (BSA), 0.1% Triton X-100, and 0.1% sodium azide). Coverslips were then washed two times in TBS for 5 minutes each prior to the addition of primary antibodies. Primary antibodies were diluted in Abdil. For KIF18A localization analyses the following primary antibodies were used at the indicated dilutions: rat anti-α-tubulin 1:500 (MAB1864; Sigma Aldrich), rabbit anti-KIF18A 1:100 (A301-080A; Bethyl), and mouse anti-Hec1 1:500 (GTX70268; GeneTex). All mCherry images for KIF18A localization analyses are direct mCherry fluorescence. For KIF18A localization analyses the following secondary antibodies were used at 1:500 dilution: Goat anti-Rabbit IgG conjugated to Alexa Fluor 488 (A11034; Invitrogen), Goat anti-Mouse IgG conjugated to Alexa Fluor 405 (A31553; Invitrogen), and Goat anti-Rat IgG conjugated to Alexa Fluor 647 (A21247, Invitrogen). For spindle length and chromosome alignment analyses, the following primary antibodies were used at the indicated dilutions: mouse anti-γ-tubulin 1:500 (T5326; Sigma Aldrich), rabbit anti-mCherry 1:500 (ab167453; Abcam), human anti-centromere antibody (ACA) 1:250 (15-235; Antibodies Inc.). All primary antibodies were incubated for 1 hour at room temperature with the exception of the human ACA antibody, which was incubated at 4°C overnight. For spindle length and chromosome alignment analyses the following secondary antibodies were used at 1:500 dilution: Goat anti-Human IgG conjugated to Alexa Fluor 488 (A11013; Invitrogen), Goat anti-Mouse IgG conjugated to Alexa Fluor 647 (A21236; Invitrogen), and Goat anti-Rabbit IgG conjugated to Alexa Fluor 594 (A11037; Invitrogen). Coverslips were washed two times in TBS for 5 minutes each between primary and secondary antibody incubations. Coverslips were washed three times in TBS for 5 minutes each prior to mounting coverslips with Prolong Gold anti-fade mounting medium with DAPI (spindle length and chromosome alignment analyses) (P36935, Invitrogen) or Prolong Gold anti-fade mounting medium without DAPI (KIF18A localization analyses) (P36934, Invitrogen). Coverslips were imaged on a Ti-E inverted microscope (Nikon Instruments) using a Plan Apo λ 60x 1.42 NA objective, environmental chamber at 37°C, a Clara cooled charge-coupled device (CCD) camera (Andor), and Nikon Elements Software (Nikon Instruments).

### Chromosome Alignment Analysis

Cells expressing mCherry or indicated mCherry-KIFBP constructs were fixed and stained for mCherry, γ-tubulin, and ACA as described above. As described previously^28,39^, single focal plane images with both spindle poles in focus were acquired. A boxed region of interest with a fixed height and width defined by the length of the spindle was used to measure the distribution of ACA-labeled kinetochore fluorescence using the Plot Profile command in Fiji. The ACA signal intensity was normalized internally to its highest value and plotted as a function of distance along the pole-to-pole axis. These plots were then fitted to a Gaussian curve and the FWHM for the Gaussian fit as well as the spindle length are reported for each cell analyzed. Mean and standard deviations are reported from a minimum of three independent experiments for each construct. The following cell numbers were analyzed for the indicated mCherry and mCherry-KIFBP constructs: (1) mCherry (control) = 132 cells, (2) mCherry-KIFBP-WT = 165 cells, (3) mCherry-KIFBP-L1^m^ = 102 cells, (4) mCherry-KIFBP-L14^m^ = 99 cells, (5) mCherry-KIFBP-HP9b^m^ = 89 cells.

### KIF18A Line Scan Analysis

Cells expressing mCherry or indicated mCherry-KIFBP constructs were fixed and stained for endogenous KIF18A, α-tubulin, and Hec1 as described above. Cells were imaged with 0.2 μm z-stacks throughout the entire cell. Within these z-sections, 2 μm line scans were manually drawn in Fiji for individual kinetochore microtubules (1-3 line scans per cell) and the profile intensities along those lines were measured and recorded for the KIF18A, α-tubulin, and Hec1 channels. Each of these profile intensities for KIF18A, α-tubulin, and Hec1 were normalized internally to its highest value. These normalized line scans were then aligned by peak Hec1 intensity and averaged for each pixel distance. Mean and standard deviations are reported from a minimum of three independent experiments for each construct. The following cell numbers and line scans were analyzed for the indicated mCherry and mCherry-KIFBP constructs: (1) mCherry (control) = 40 cells (64 lines), (2) mCherry-KIFBP-WT = 34 cells (64 lines), (3) mCherry-KIFBP-L1^m^ = 34 cells (64 lines), (4) mCherry-KIFBP-L14^m^ = 32 cells (68 lines), (5) mCherry-KIFBP-HP9b^m^ = 33 cells (63 lines).

### Molecular Dynamics Simulations and Analysis

The structures of KIF5C, KIF15, KIF18A, and KIF11 bound to ADP and Mg^2+^ were taken from PDB structures 1BG2^74^, 4BN2^36^, 3LRE^71^, and 1II6^75^, respectively. The missing residues of KIF5C were filled in as previously described^76^. For all other proteins, I-TASSER was used to fill in the gaps of the remaining structures using the PDBs as primary template^77–79^. AmberTools was then used to prepare all systems for simulation^80^. Each system was solvated with a box of TIP3P water molecules with 10-Å padding around the protein. Na^+^ and Cl^-^ were added to both neutralize the systems and set the ionic concentration to 50 mM. NAMD was used to carry out the MD simulations with the amber ff19SB force field^81,82^. Force field parameters for the ADP nucleotide were obtained from the AMBER parameter database^83^. Following minimization, heating and equilibration, the systems were simulated at 300 K and 1 atm of pressure in an NpT ensemble. To allow for 2-fs time steps, bonded hydrogens were fixed. For long-range electrostatics, Particle Mesh Ewald was employed with a 10-Å cutoff and 8.5-Å switch distance for van der Waals interactions^84^. MD simulations were completed in 100 ns replicates starting from random velocities for a total simulation time of 500 ns for KIF5C, KIF11 and KIF18A, and 600 ns for KIF15.

All analysis was carried out using the Bio3d package (v 2.4.1) in R^85^. We first aligned our 4 kinesins of interest and then restricted analysis to the amino acids that appeared in all the motors. In order to focus on large-scale rearrangements in the motor head and remove the noise from fluctuating loops, we next restricted analysis to amino acids in stable secondary structures, leaving us with 154 amino acid positions in each protein. We performed Principal Component Analysis (PCA) using KIF15 as the reference structure. The remaining three kinesins simulations were then projected onto this KIF15 PCA space for direct comparison with each other and the cryo-EM structure.

