## Supplementary Material for "Kinesin-binding protein remodels the kinesin motor to prevent microtubule-binding"

### TABLES

#### Table 1. Cryo-EM data collection, analysis, and validation statistics for KIFBP (full).

| | **Structure: KIFBP (full)** (EMDB-xxxx) (PDB xxxx) | | |
|---|---|---|---|
| **Data collection** | | | |
| Grids | UltrAuFoil | UltrAuFoil | UltrAuFoil |
| Vitrification method | FEI Vitrobot | FEI Vitrobot | FEI Vitrobot |
| Microscope | Glacios | Glacios | Glacios |
| Session name | 20jul13c | 20jul07a | 20jul11b |
| Magnification | 45000X | 45000X | 45000x |
| Voltage (kV) | 200 | 200 | 200 |
| Stage tilt (°) | 0 | 0 | 0 |
| Detector | K2 Summit | K2 Summit | K2 Summit |
| Recording mode | Counting | Counting | Counting |
| Total electron exposure ($e^{-}/Å^2$) | 58.5 | 68.4 | 65.0 |
| Number of frames | 45 | 40 | 35 |
| Defocus range (μm) | 0.8 – 2.0 | 0.8 – 2.0 | 0.8-2.0 |
| Pixel size (Å) | 0.98 | 0.98 | 0.98 |
| **Data processing** | | | |
| Number of micrographs | 5,349 | 1,596 | 4,144 |
| Initial particle images (no.) | 1,121,348 | 403,869 | 913,455 |
| Final particle images (no.) | 115,464 | 32,422 | 43,210 |
| Final particles in refinement | 154,176 | | |
| Symmetry | C1 | | |
| Map resolution (Å) | 4.6 | | |
| **Refinement** | | | |
| Initial model used (PDB code) | N/A | | |
| Cryo-efficiency (cryoEF) | 0.7 | | |
| Model resolution (Å) | 4.6 | | |
| FSC threshold | 0.143 | | |
| Map sharpening *B* factor ($Å^2$) | -50 | | |
| Model composition | | | |
| Non-hydrogen atoms | 4332 | | |
| Protein residues | 527 | | |
| Ligands | 0 | | |
| *B* factors ($Å^2$) | N/A | | |
| Protein | | | |
| Ligand | | | |
| R.m.s. deviations | N/A | | |
| Bond lengths (Å) | | | |
| Bond angles (°) | | | |
| Validation | N/A | | |
| MolProbity score | | | |
| Clashscore | | | |
| Poor rotamers (%) | | | |
| Ramachandran plot | N/A | | |
| Favored (%) | | | |
| Allowed (%) | | | |
| Disallowed (%) | | | |

**Table 2. Cryo-EM data collection, analysis, and validation statistics for KIFBP (core).**

| | **Structure: KIFBP (core)**<br>(EMDB-xxxx)<br>(PDB xxxx) |
|---|---|
| **Data collection** | |
| Grids | UltrAuFoil |
| Vitrification method | FEI Vitrobot |
| Microscope | Glacios |
| Session name | 20jul11b |
| Magnification | 45000X |
| Voltage (kV) | 200 |
| Stage tilt (°) | 0 |
| Detector | K2 Summit |
| Recording mode | Counting |
| Total electron exposure ($e^{-}$/Å$^2$) | 65 |
| Number of frames | 35 |
| Defocus range (μm) | 0.8 – 2.0 |
| Pixel size (Å) | 0.98 |
| **Data processing** | |
| Number of micrographs | 4,144 |
| Initial particle images (no.) | 913,455 |
| Final particles in refinement | 128,190 |
| Symmetry | C1 |
| Map resolution (Å) | 3.8 |
| **Refinement** | |
| Cryo-efficiency (cryoEF) | 0.5 |
| FSC model to map (0.143, 0.5) | 3.5, 4.5 |
| Initial model used (PDB code) | N/A |
| Model resolution (Å) | 3.8 |
| FSC threshold | 0.143 |
| Map sharpening $B$ factor (Å$^2$) | -50 |
| Model composition | |
| Non-hydrogen atoms | 2015 |
| Protein residues | 246 |
| Ligands | 0 |
| $B$ factors (Å$^2$) | |
| Protein | 57.76 |
| Ligand | N/A |
| R.m.s. deviations | |
| Bond lengths (Å) | 0.008 |
| Bond angles (°) | 0.942 |
| Validation | |
| MolProbity score | 1.65 |
| Clashscore | 3.25 |
| Poor rotamers (%) | 2.88 |
| Ramachandran plot | |
| Favored (%) | 96.90 |
| Allowed (%) | 3.10 |
| Disallowed (%) | 0 |

**Table 3. Cryo-EM data collection, analysis, and validation statistics for KIFBP:KIF15.**

| | **Structure: KIFBP:KIF15** (EMDB-xxxx) (PDB xxxx) | | | |
|---|---|---|---|---|
| **Data collection** | | | | |
| Grids | UltrAuFoil | UltrAuFoil | UltrAuFoil | UltrAuFoil |
| Vitrification method | FEI Vitrobot | FEI Vitrobot | FEI Vitrobot | FEI Vitrobot |
| Microscope | Glacios | Glacios | Glacios | Glacios |
| Session name | 20aug23f | 20aug30b | 20sep17h | 20sep25f |
| Magnification | 45000X | 45000X | 45000X | 45000X |
| Voltage (kV) | 200 | 200 | 200 | 200 |
| Stage tilt (°) | 0 | 0 | 0 | 0 |
| Detector | K2 Summit | K2 Summit | K2 Summit | K2 Summit |
| Recording mode | Counting | Counting | Counting | Counting |
| Total electron exposure ($e^{-}$/Å$^2$) | 83.6 | 60.1 | 61.25 | 60.76 |
| Number of frames | 40 | 30 | 35 | 35 |
| Defocus range (μm) | 0.8 – 2.0 | 0.8 – 2.0 | 0.8-2.0 | 0.8-2.0 |
| Pixel size (Å) | 0.98 | 0.98 | 0.98 | 0.98 |
| **Data processing** | | | | |
| Number of micrographs | 1,007 | 972 | 1,247 | 2,958 |
| Initial particle images (no.) | 324,933 | 282,365 | 394,676 | 1,031,191 |
| Final particle images (no.) | 25,182 | 32,231 | 41,674 | 90,897 |
| Final particles in refinement | 101,698 | | | |
| Symmetry | C1 | | | |
| Map resolution (Å) | 4.8 | | | |
| **Refinement** | | | | |
| Initial model used (PDB code) | 4BN2 | | | |
| Cryo-efficiency (cryoEF) | 0.58 | | | |
| Model resolution (Å) | 4.8 | | | |
| FSC threshold | 0.143 | | | |
| FSC model to map (0.143, 0.5) | 4.8, 7.1 | | | |
| Map sharpening *B* factor (Å$^2$) | -149 | | | |
| Model composition | | | | |
| Non-hydrogen atoms | 5699 | | | |
| Protein residues | 725 | | | |
| Ligands | 0 | | | |
| *B* factors (Å$^2$) | | | | |
| Protein | 199 | | | |
| Ligand | 0 | | | |
| R.m.s. deviations | | | | |
| Bond lengths (Å) | 0.015 | | | |
| Bond angles (°) | 1.382 | | | |
| Validation | | | | |
| MolProbity score | 1.68 | | | |
| Clashscore | 4.15 | | | |
| Poor rotamers (%) | 0 | | | |
| Ramachandran plot | | | | |
| Favored (%) | 92.08 | | | |
| Allowed (%) | 6.22 | | | |
| Disallowed (%) | 1.70 | | | |

**Table 4. Cryo-EM data collection, analysis, and validation statistics for KIFBP:KIF18A.**

| | **Structure: KIFBP:KIF18A** (EMDB-xxxx) (PDB xxxx) | | |
|---|---|---|---|
| **Data collection** | | | |
| Grids | UltrAuFoil | UltrAuFoil | UltrAuFoil |
| Vitrification method | FEI Vitrobot | FEI Vitrobot | FEI Vitrobot |
| Microscope | Talos Arctica | Talos Arctica | Talos Arctica |
| Session name | 20oct06g | 20nov05c | 21jan22b |
| Magnification | 45000X | 45000X | 45000x |
| Voltage (kV) | 200 | 200 | 200 |
| Stage tilt (°) | 0 | 0 | 0 |
| Detector | K2 Summit | K2 Summit | K2 Summit |
| Recording mode | Counting | Counting | Counting |
| Total electron exposure ($e^{-}/Å^2$) | 57.0 | 51.8 | 61.2 |
| Number of frames | 35 | 35 | 40 |
| Defocus range (µm) | 0.8 – 2.0 | 0.8 – 2.0 | 0.8-2.0 |
| Pixel size (Å) | 0.91 | 0.91 | 0.91 |
| **Data processing** | | | |
| Number of micrographs | 532 | 571 | 3,566 |
| Initial particle images (no.) | 71,529 | 94,716 | 638,186 |
| Final particle images (no.) | 44,395 | | 111,764 |
| Final particles in refinement | 54,801 | | |
| Symmetry | C1 | | |
| Map resolution (Å) | 4.9 | | |
| **Refinement** | | | |
| Initial model used (PDB code) | 3LRE | | |
| Cryo-efficiency (cryoEF) | 0.67 | | |
| Model resolution (Å) | 4.9 | | |
| FSC threshold | 0.143 | | |
| Map sharpening *B* factor ($Å^2$) | -163.5 | | |
| Model composition | N/A | | |
| Non-hydrogen atoms | | | |
| Protein residues | | | |
| Ligands | | | |
| *B* factors ($Å^2$) | N/A | | |
| Protein | | | |
| Ligand | | | |
| R.m.s. deviations | N/A | | |
| Bond lengths (Å) | | | |
| Bond angles (°) | | | |
| Validation | N/A | | |
| MolProbity score | | | |
| Clashscore | | | |
| Poor rotamers (%) | | | |
| Ramachandran plot | N/A | | |
| Favored (%) | | | |
| Allowed (%) | | | |
| Disallowed (%) | | | |

**Table 5. High-confidence crosslinks between KIFBP and KIF15.**

A summary of all high-confidence crosslinks identified between KIFBP and the KIF15 motor domain using mass spectrometry and the lysine-targeting crosslinker BS3. The position of KIFBP and KIF15 residues of each crosslink are shown.

| KIFBP Residue | KIF15 Residue |
|---|---|
| 26 | 273 |
| | 283 |
| | 319 |
| | 361 |
| 30 | 273 |
| | 283 |
| | 319 |
| | 361 |
| | 364 |
| | 366 |
| 36 | 273 |
| | 283 |
| 205 | 273 |
| | 283 |
| 307 | 283 |
| | 319 |
| 350 | 364 |
| 556 | 361 |
| | 366 |
| 564 | 283 |
| | 364 |
| 610 | 319 |
| | 361 |
| | 364 |
| | 366 |
| 617 | 361 |
| | 364 |

### SUPPLEMENTAL FIGURES

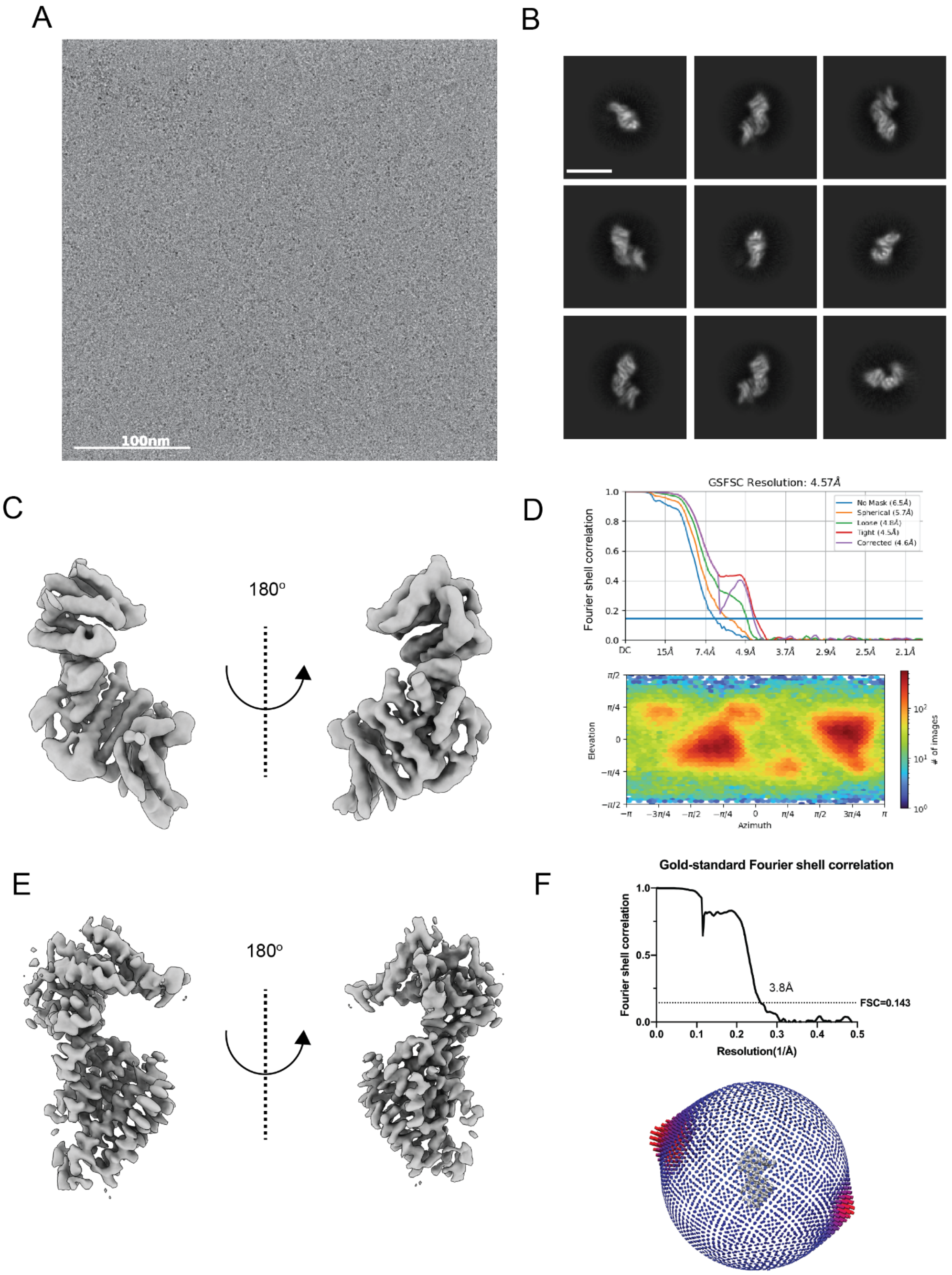

**Figure 1 - Supplement 1. Cryo-EM structures of KIFBP.**

(A) Representative cryo-EM micrograph. (B) Representative 2D class averages. Scale bar is 100Å. Full KIFBP reconstruction (C), FSC curves and euler angle distribution (D). KIFBP core overview (E), FSC curves and Euler angle distribution (F).

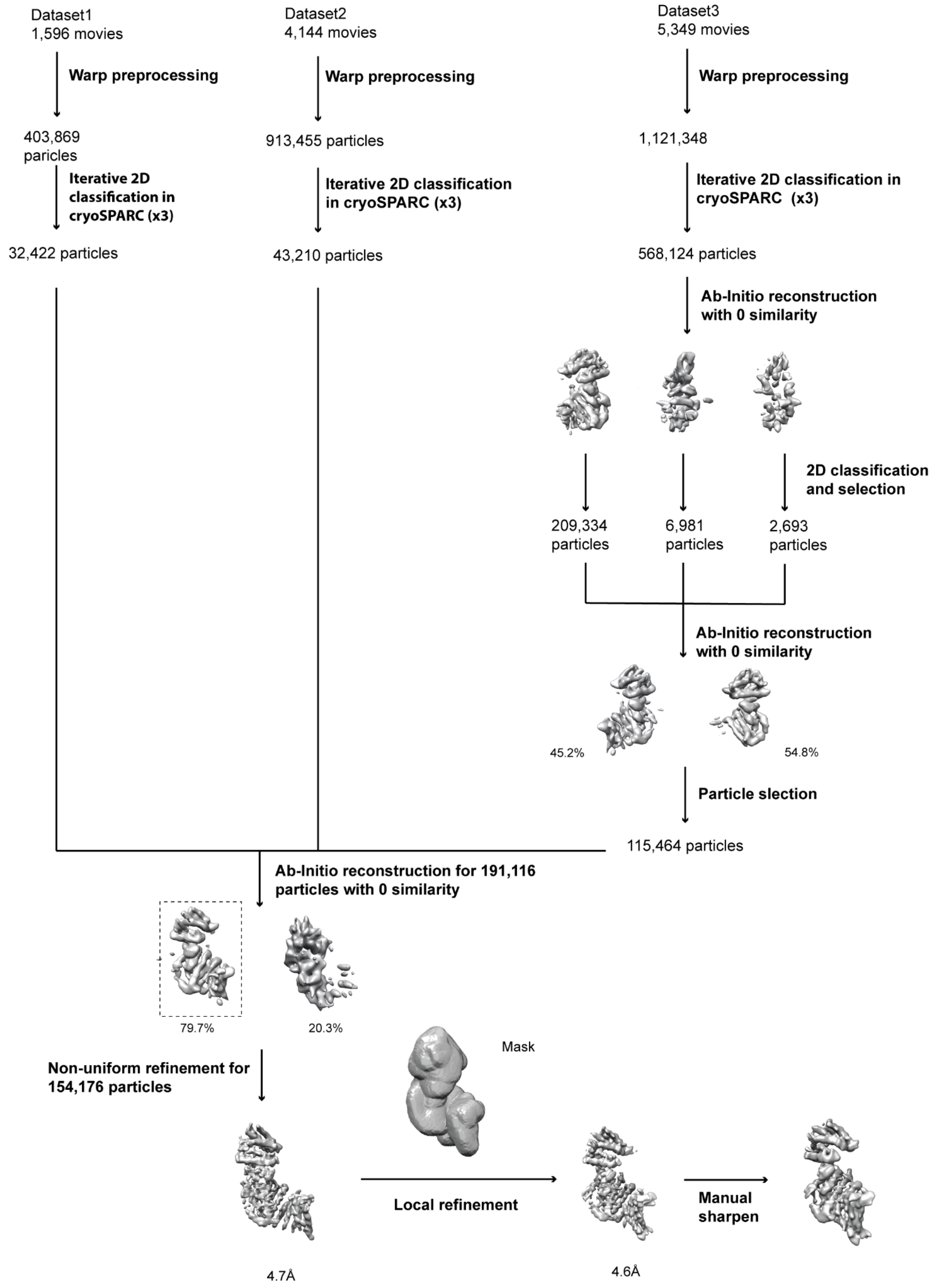

**Figure 1 - Supplement 2. Cryo-EM processing tree for 4.6Å full KIFBP reconstruction.**
Overview of data processing strategy for full KIFBP.

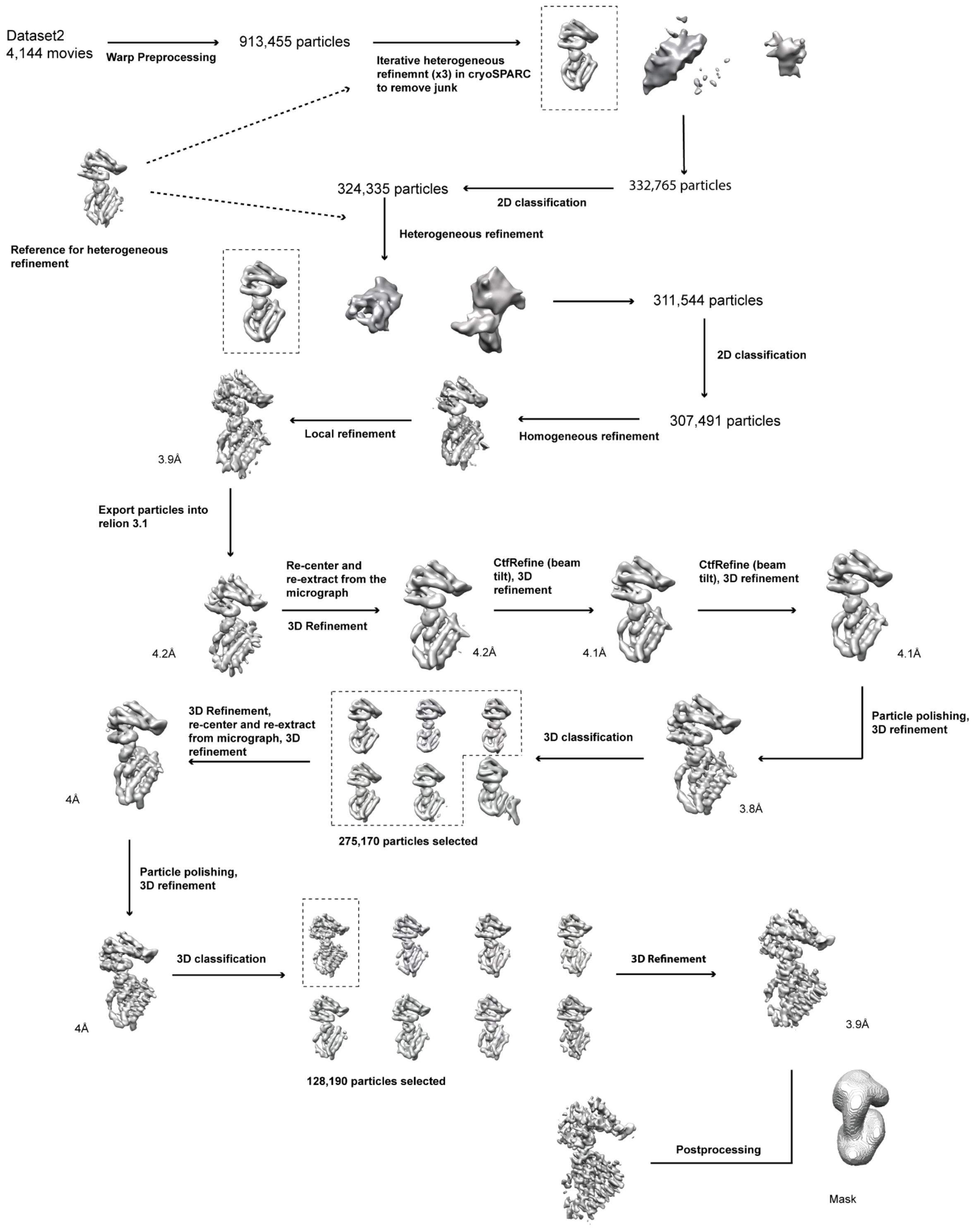

**Figure 1 - Supplement 3. Cryo-EM processing tree for 3.8Å core KIFBP reconstruction.**
Overview of data processing strategy for core KIFBP reconstruction.

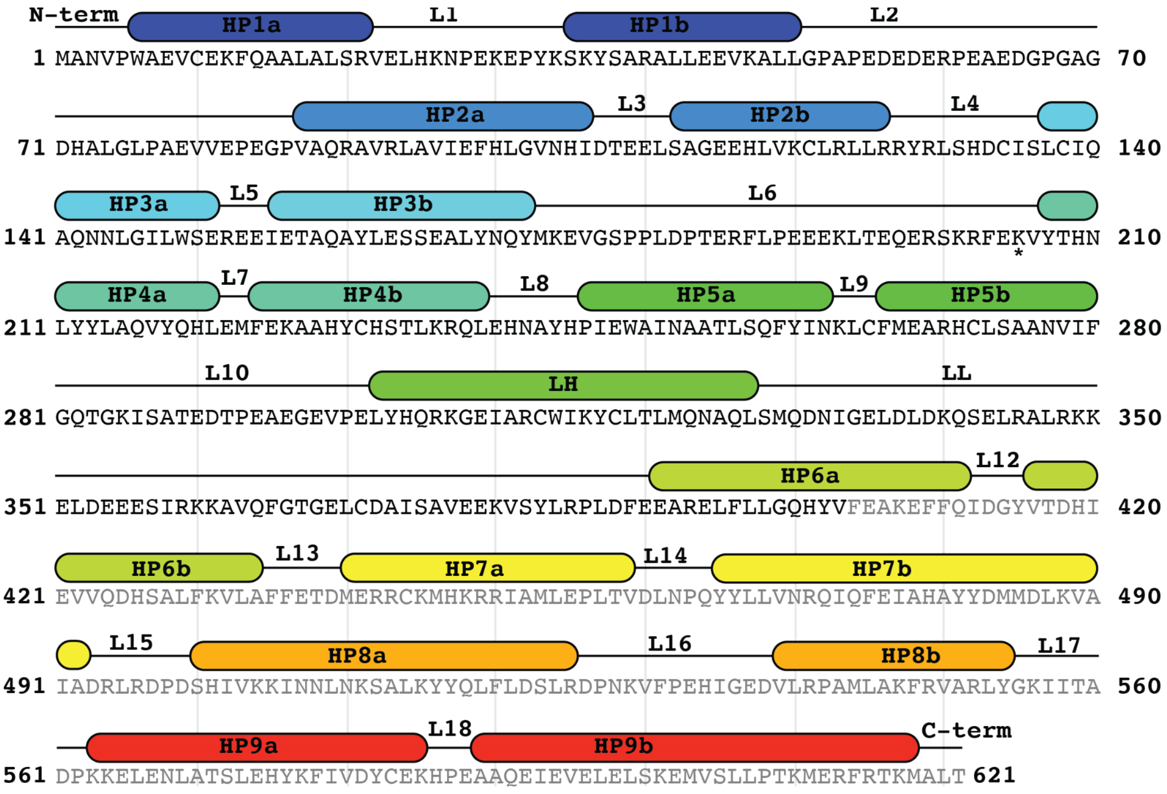

**Figure 1 - Supplement 4. Annotated KIFBP primary sequence using secondary structure information from the atomic model.**

Residues 5-403 were built *de novo* into the KIFBP core reconstruction at 3.8Å (black letters), whereas 404-621 were modeled into the 4.6Å reconstruction (gray letters).

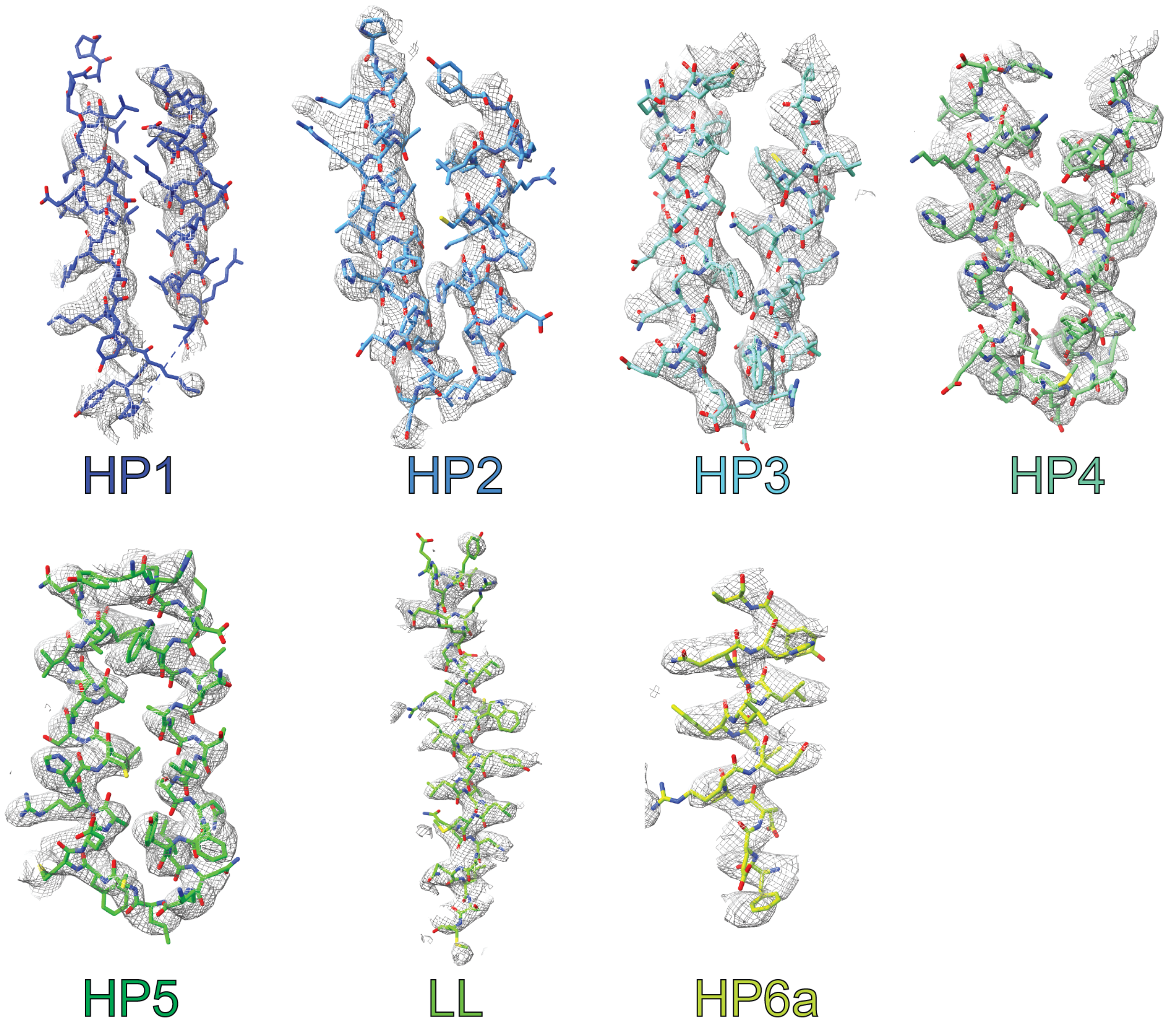

**Figure 1 - Supplement 5. Segmented density for 3.8Å core KIFBP reconstruction and atomic model.**

Helical pairs from core KIFBP reconstruction.

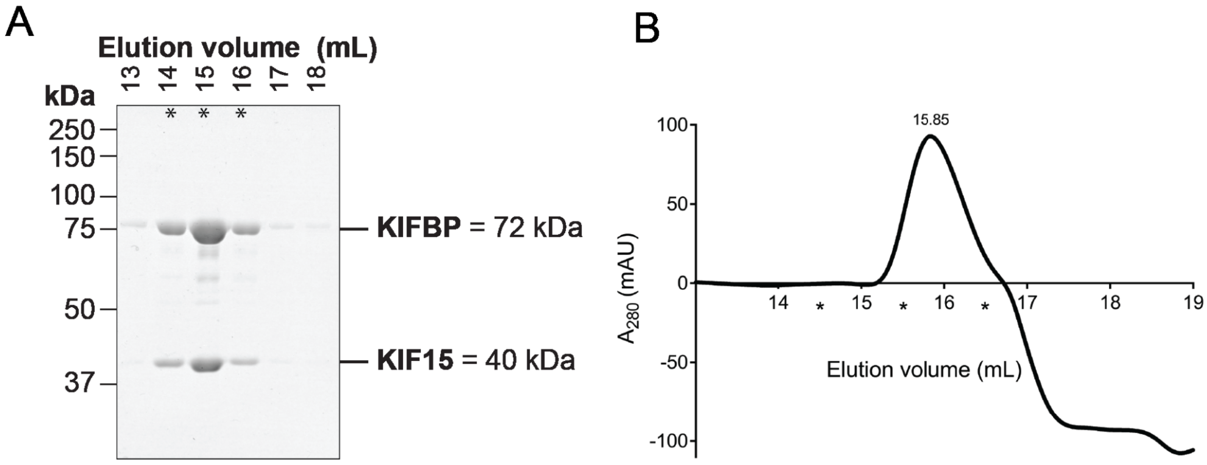

**Figure 2 - Supplement 1. Size exclusion chromatography of KIFBP:KIF15.**
KIFBP and KIF15 were purified, combined, and run over the Superose 6 column as described in methods. (A) Representative Coomassie-stained SDS-PAGE of peak fractions from elution profile shown in (B). Fractions 14, 15, and 16 were combined and used in subsequent cryo-EM experiments (indicated by asterisks). The molecular weight of each protein is indicated in kilodaltons.

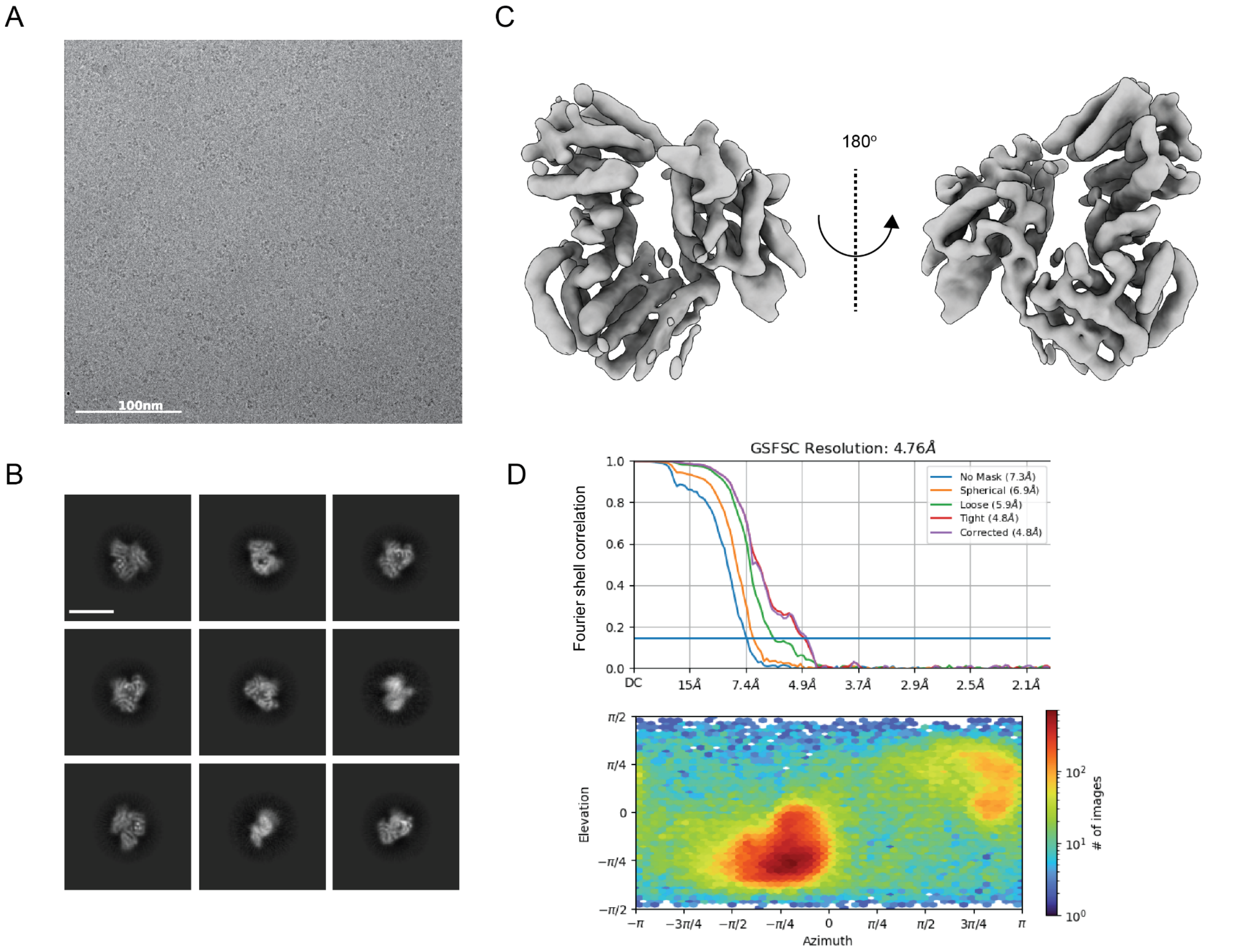

**Figure 2 - Supplement 2. Cryo-EM structure of KIFBP:KIF15.**
(A) Representative micrograph for KIFBP:KIF15. (B) Representative 2D class averages. The scale bar is 100Å. (C) Reconstruction overview. (D) FSC curves and Euler angle distribution.

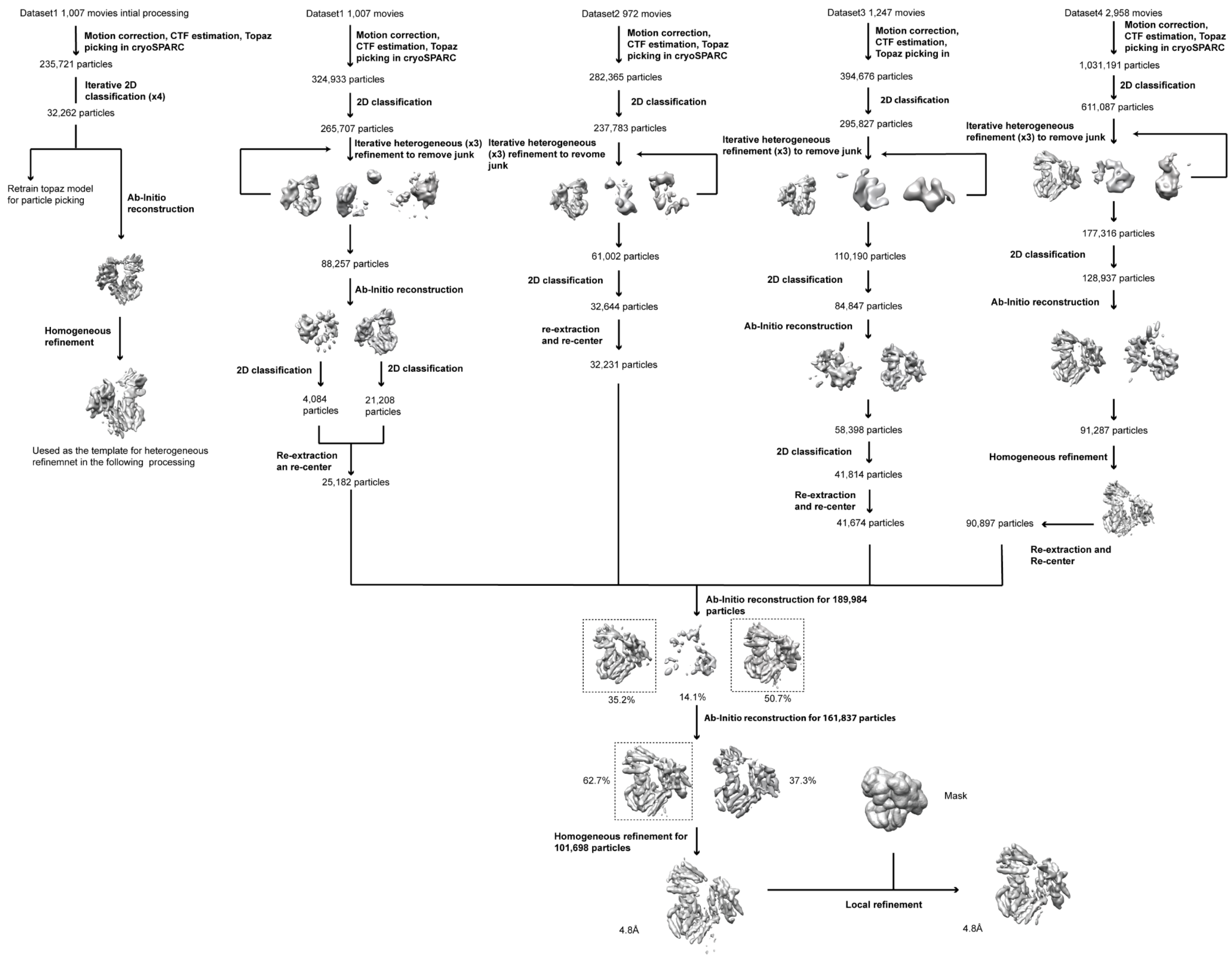

**Figure 2 - Supplement 3. Cryo-EM processing tree for KIFBP:KIF15.**
Overview of processing steps for KIFBP:KIF15.

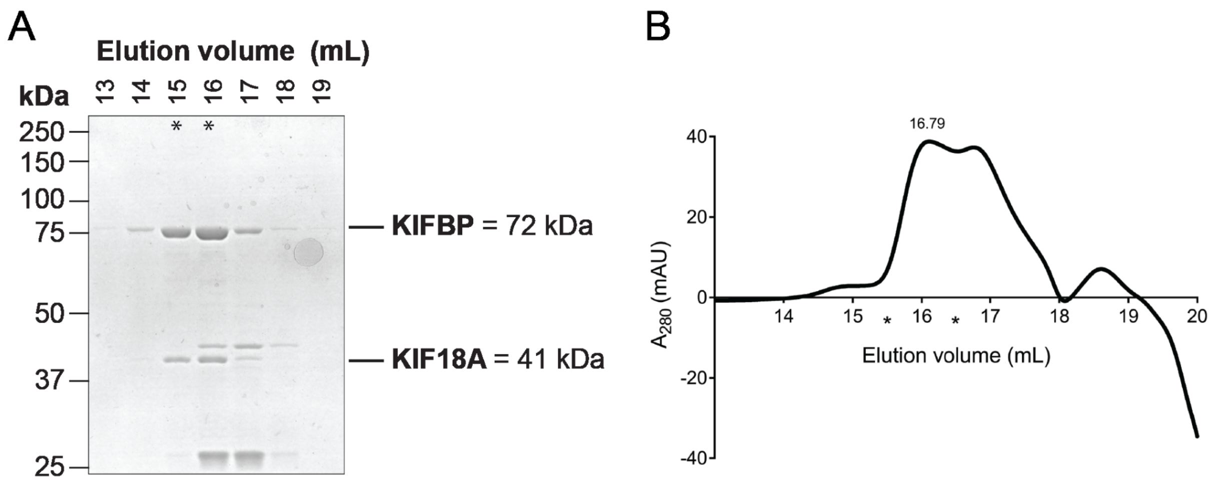

**Figure 5 - Supplement 1. Size exclusion chromatography of KIFBP:KIF18A.**
KIFBP and KIF18A were purified, combined, and run over the Superose 6 column as described in methods. (A) Representative Coomassie-stained SDS-PAGE of peak fractions from elution profile shown in (B). Fractions 15 and 16 were combined and used in subsequent cryo-EM experiments (indicated by asterisks). The molecular weight of each protein is indicated in kilodaltons.

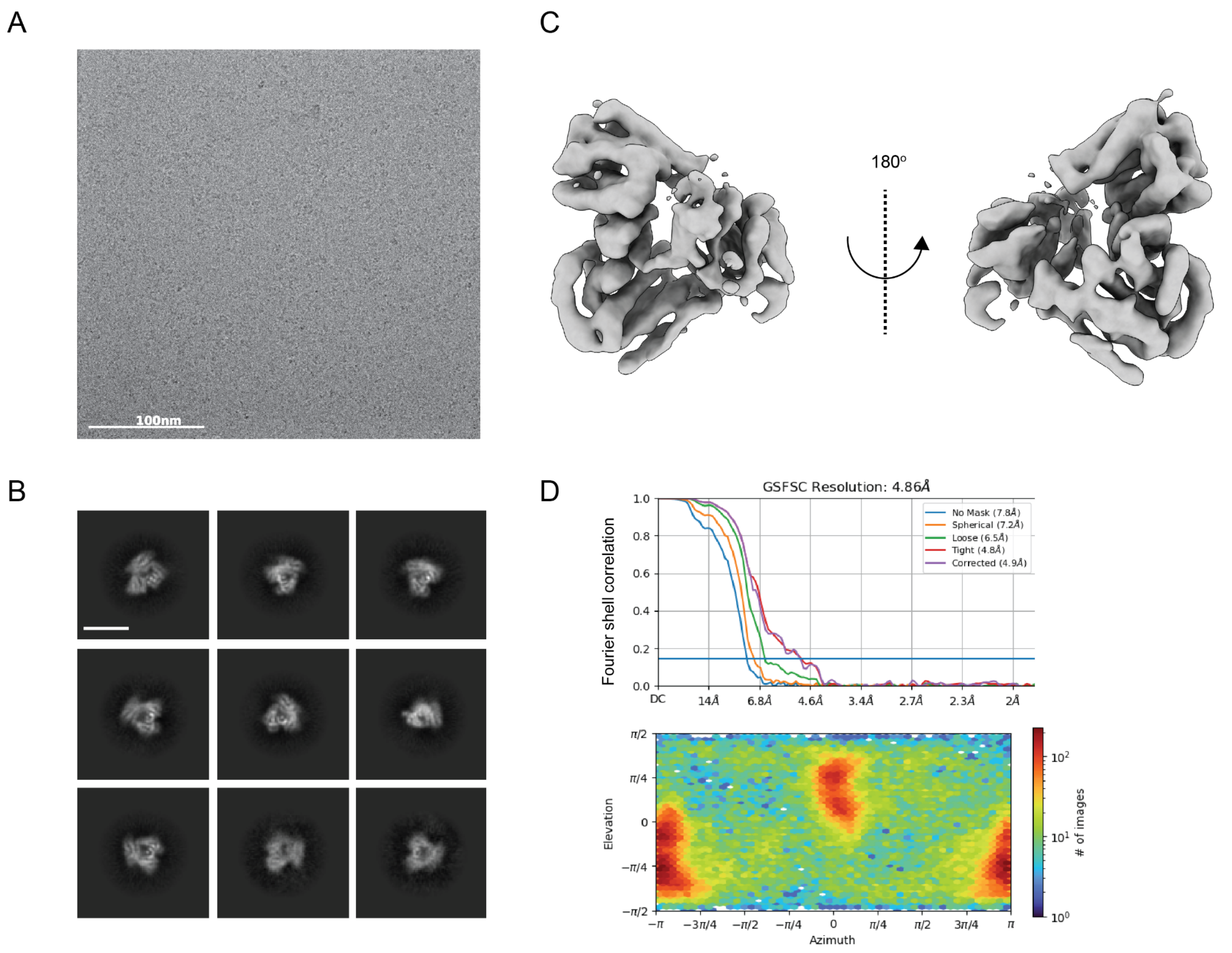

**Figure 5 - Supplement 2. Cryo-EM structure of KIFBP:KIF18A.**

(A) Representative micrograph for KIFBP:KIF18A. (B) Representative 2D class averages. Scale bar is 100Å. (C) Reconstruction overview. (D) FSC curves and Euler angle distribution.

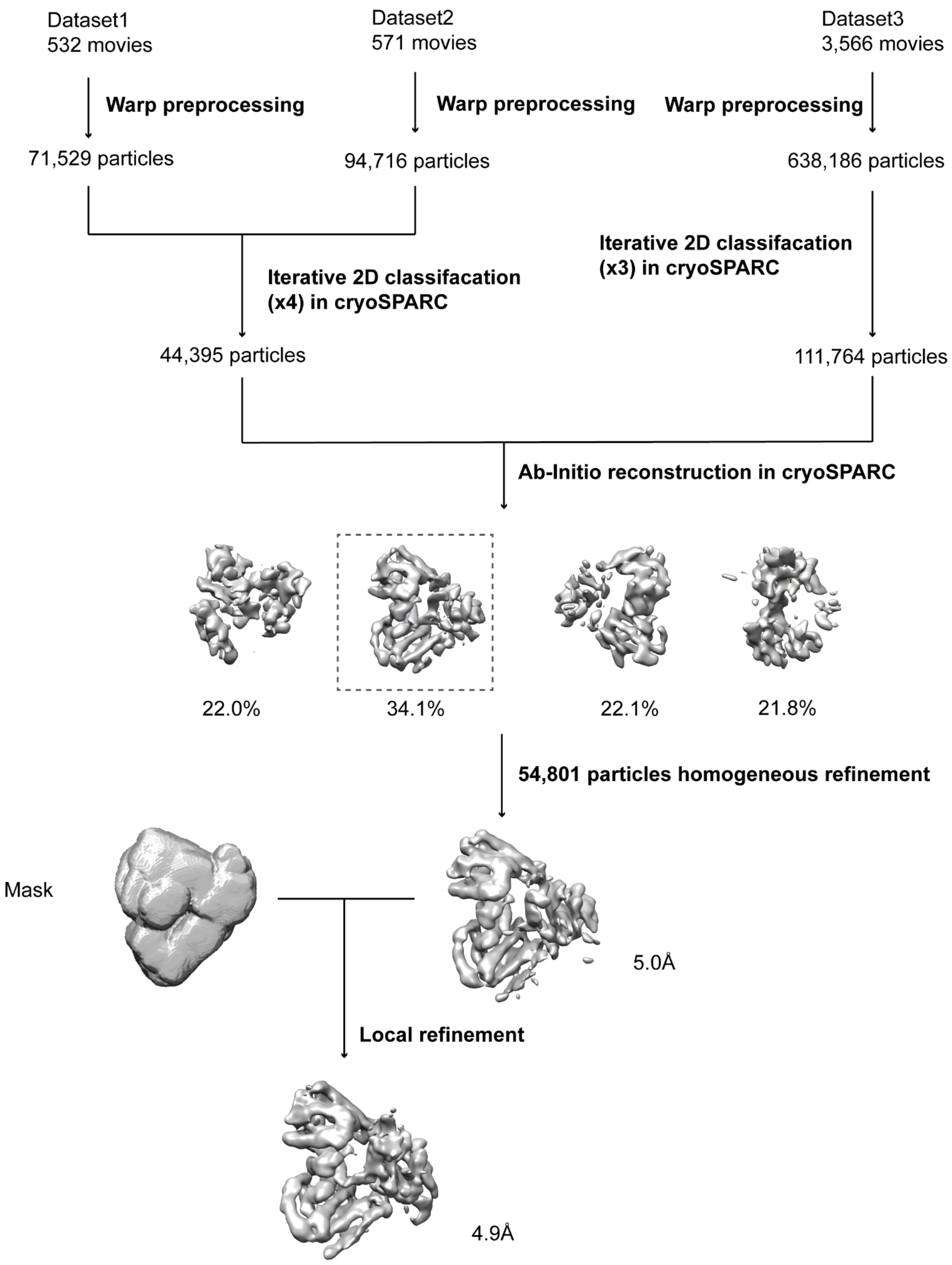

**Figure 5 - Supplement 3. Cryo-EM processing tree for KIFBP:KIF18A.**
Overview of analysis strategy for KIFBP:KIF18A.

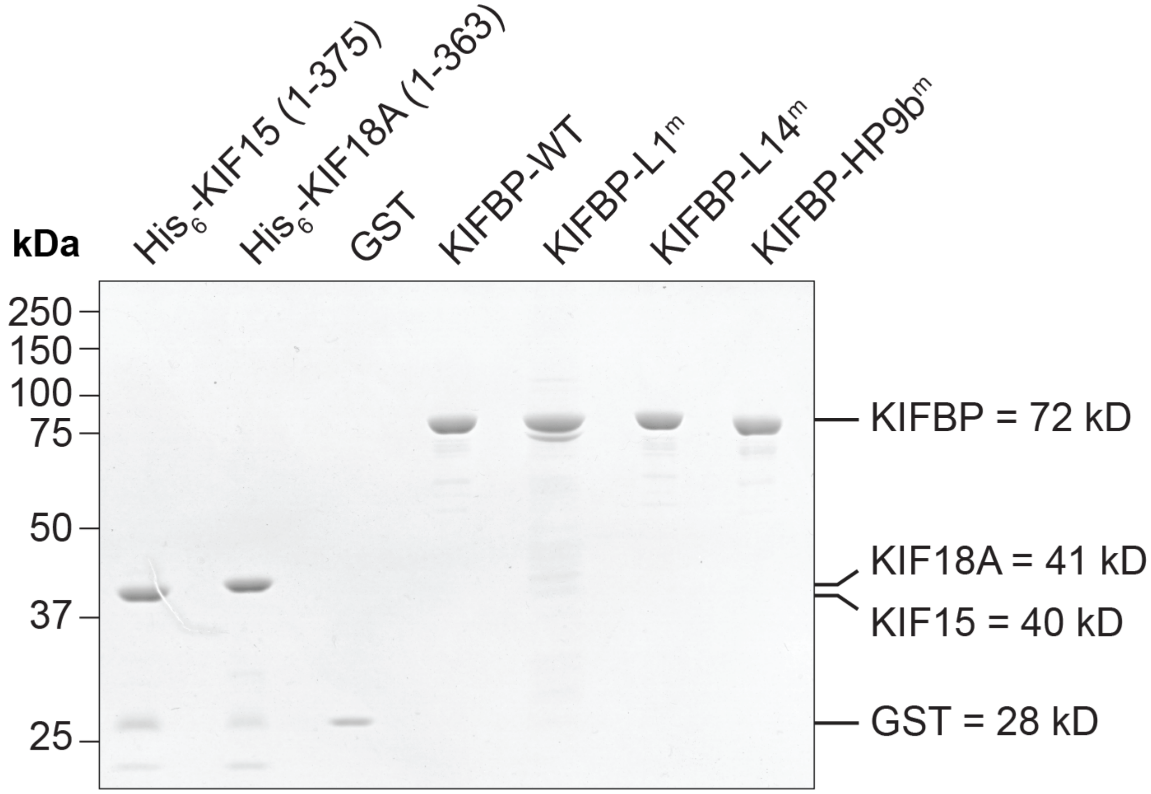

**Figure 6 - Supplement 1. Input of individual proteins used in the pull-down binding assay.** Representative Coomassie gel of 1 µg of each protein used in the pull-down assays in Figure 6C & D.

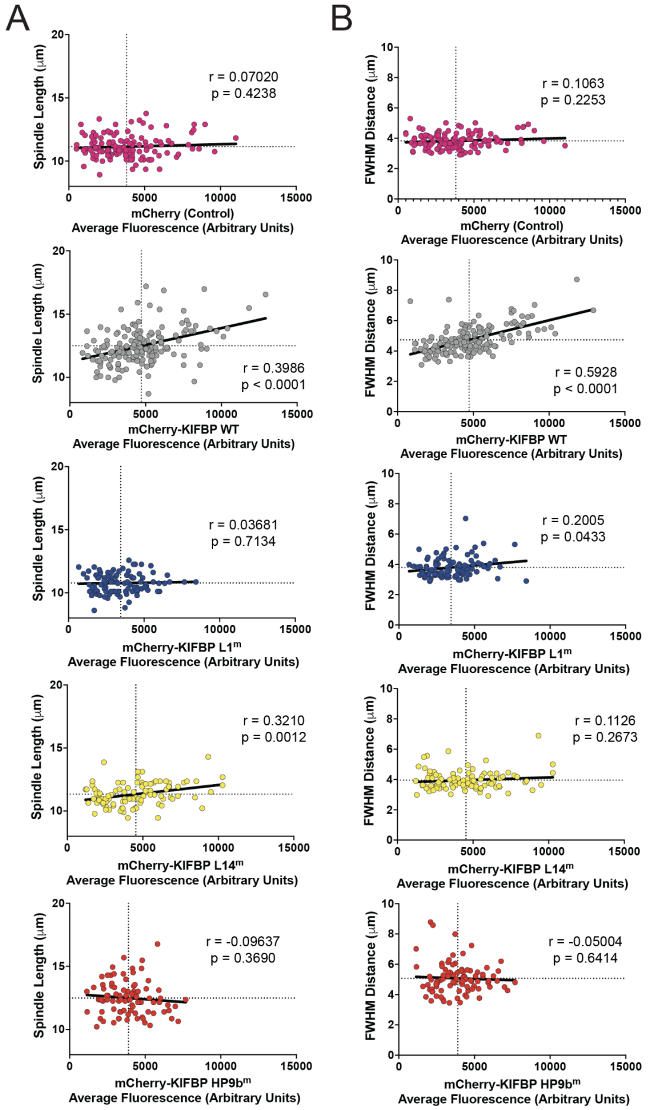

**Figure 7 - Supplement 1. Mitotic effects of mCherry-KIFBP-WT scale with expression level.**

(A) Plots of Spindle Length versus Average mCherry Fluorescence for HeLa Kyoto cells overexpressing mCherry or indicated mCherry-KIFBP construct. Each dot represents a single cell. Data presented from a minimum of three independent experiments. Dotted lines represent the mean value for Spindle Length or Average mCherry Fluorescence. Solid line is a linear regression showing the trend of the data. The Pearson's correlation coefficient (r) and two-tailed p-value with 95% confidence interval are shown for each plot. (B) Plots of Full-Width at Half Maximum (FWHM) Distance versus Average mCherry Fluorescence for HeLa Kyoto cells overexpressing mCherry or indicated mCherry-KIFBP construct. Each dot represents a single cell. Data presented from a minimum of three independent experiments. Dotted lines represent the mean value for FWHM distance or Average mCherry Fluorescence. Solid line is a linear regression showing the trend of the data. The Pearson's correlation coefficient (r) and two-tailed p-value with 95% confidence interval are shown for each plot.
